# The substrate quality of CK2 target sites has a determinant role on their function and evolution

**DOI:** 10.1101/2023.07.03.547533

**Authors:** David Bradley, Chantal Garand, Hugo Belda, Isabelle Gagnon-Arsenault, Moritz Treeck, Sabine Elowe, Christian R Landry

## Abstract

Most biological processes are regulated by peptide-recognition modules (PRMs) that bind to short linear motifs (SLiMs). Such interactions are rapidly reversible and often occur at low affinity. The protein kinase domain represents one such binding module, and known substrates may have full or only partial matches to the kinase recognition motif, a property known as ‘substrate quality’. However, it is not yet clear whether differences in substrate quality represent neutral variation along the phosphosite sequence or if these differences have functional consequences that are subject to selection. We explore this question in detail for the acidophilic kinase CK2. CK2 is well-characterised, clinically important, and a fundamental enzyme for many aspects of cell biology. We show that optimal CK2 sites are phosphorylated at maximal stoichiometries and found in many conditions whereas minimal substrates are phosphorylated at lower stoichiometries, are more dynamic during the cell cycle, and have regulatory functions. Optimal CK2 sites also tend to be older and more conserved than minimal sites, and evolutionary simulations indicate that the substrate quality of CK2 phosphosites is often tuned by selection. For intermediate target sites, increases or decreases to substrate quality may be deleterious, which we demonstrate experimentally for a CK2 substrate at the kinetochore. The results together suggest that minimal and optimal phosphosites are strongly differentiated in terms of their functional and evolutionary properties.

## Introduction

The specificity of protein kinases is determined in part by amino acid residues in the substrate-binding pocket (Ubersax and Ferrell 2007; Creixell et al. 2015; C. J. Miller and Turk 2018; Bradley et al. 2021). These define a degenerate phosphorylation motif of the kinase that denotes favoured substrate residues at the active site. Substrates for hundreds of human kinases have now been determined from *in vitro* and *in vivo* assays (Hornbeck et al. 2015, 2019; Licata et al. 2020; Bachman, Gyori, and Sorger 2022). However, phosphosites may display a range of motif matching scores for their cognate kinase, with many having just the minimal match to the kinase preferred sequence (Oliveira et al. 2016; Needham et al. 2019). The existence of such ‘minimal’ phosphosites implies evolutionary constraint against the optimisation of kinase-substrate networks, for which there are a number of possible reasons.

Most trivially, a small number could be explained by literature curation errors from databases of kinase-substrate relationships (KSRs) such as PhosphoSitePlus and Signor (Hornbeck et al. 2015; Needham et al. 2019; Licata et al. 2020; Invergo 2022; Bachman, Gyori, and Sorger 2022). A potentially larger number of motifs are minimal in the primary structure but can be reconstituted as full motifs when the target sequence folds in three dimensions to accommodate the substrate-binding pockets of the kinase (Chihao Wang et al. 2012; Duarte et al. 2014; Oliveira et al. 2016).

Minimal substrates may also be explained by constraint in cellular networks. Kinases can be considered as ‘writer’ enzymes for phosphosites and by analogy phosphatases and phosphate-binding domains serve as ‘erasers’ and ‘readers’ (respectively) (Pincus et al. 2008; Lim and Pawson 2010; Beltrao et al. 2013). Phosphate-binding domains (SH2, PTB, BRCT, 14-3-3, WW, FHA, etc) and phosphatase domains have their own peptide specificities and to varying extents the evolution of the phosphosite sequence must satisfy constraints across the three domain classes simultaneously (Otte et al. 2003; Smith et al. 2006; M. L. Miller et al. 2008; Tinti et al. 2013; X. Li et al. 2013; Damle and Köhn 2019; Cantor, Shah, and Kuriyan 2018). This principle applies also to phosphosites targeted by more than one kinase, and to clustered phosphosites with overlapping flanking sequences in the primary structure (Moses, Hériché, and Durbin 2007; Schweiger and Linial 2010). An extreme example is given by hierarchical phosphorylation sites, where an initial phosphorylation ‘primes’ a secondary phosphorylation within 3-4 amino acid residues of the first phosphosite (Nishi, Shaytan, and Panchenko 2014; Cesaro and Pinna 2015; N. St-Denis et al. 2015). Finally, there are more global constraints at the level of the whole protein in terms of the stability of protein folding or protein-protein interactions, which can be impacted by the phosphosite flanking sequence depending on the phosphosite position in the tertiary structure (Nishi, Shaytan, and Panchenko 2014; Cole and Prabakaran 2020). The need to satisfy all of these constraints at the same time could explain why many phosphosites are suboptimal for their respective kinase. Such constraints also imply that most phosphosites will exist between the extremes of minimality and optimality in terms of the kinase recognition motif.

Many phosphosites identified by mass spectrometry are also likely to have no function and could instead represent off-target kinase activity (Lienhard 2008; Landry, Levy, and Michnick 2009; Nguyen Ba and Moses 2010; Levy, Michnick, and Landry 2012; Beltrao et al. 2012). For such sites, optimising the flanking sequence would confer no selective advantage and may be deleterious if the spurious phosphosite has a significant fitness cost. Even for functional phosphosites, there is a heavy mutational bias against the optimization of the phosphorylation motif; in other words, a spontaneous mutation is much more likely to decrease than increase the quality of the substrate for its cognate kinase(s) (Wagih, Reimand, and Bader 2015; Lynch and Hagner 2015; Lynch 2018). Whether an optimised phosphosite could survive the randomness of drift and mutation would therefore depend on the selection differential between the optimal and minimal genotypes of the phosphosite.

Finally, a suboptimal phosphorylation efficiency may yield a desirable output in the context of a signalling pathway or network. For example, a recent computational model of kinase-substrate pathways suggests that increasing the strength of kinase-substrate interactions leads to a dynamical response that is slower and less reversible (C.-H. Wang, Mehta, and Bashor 2018). This is consistent with the more general supposition that domain-peptide interactions are maintained at relatively weak affinities because it enables signalling outputs to be reversed amid fluctuating external stimuli (Pawson 2004; Ladbury and Arold 2012; Haslam and Shields 2012; Davey et al. 2012). An example was found in 2019 for the ZAP70-LAT kinase-substrate interaction in T-cells. The LAT Y132 phosphosite sequence is conserved across orthologues to be suboptimal as it features glycine at the −1 position (relative to the phosphoacceptor) instead of the favoured aspartate at −1; ‘optimization’ of phosphosite by mutation accelerates the kinetics of phosphorylation and leads to the maladaptive sensitisation of T-cells to self-antigens (Lo et al. 2019).

For this study we aim to explore systematically the relationship between substrate quality and phosphorylation stoichiometry, function, and evolution using computational and experimental approaches. Towards this end, we focus on the substrates of CK2, which is a highly pleiotropic kinase with over **500** known substrates in human (Meggio and Pinna 2003; Hornbeck et al. 2019). CK2 has a minimal substrate phosphorylation motif of **S/T**-x-x-**D/E** but has significant aspartate/glutamate preferences for all substrate positions from **-5** to **+5** and is depleted for positively charged **K/R** residues (Hutti et al. 2004; Chunli Wang et al. 2013; Hornbeck et al. 2019; Johnson et al. 2023). The CK2 ‘substrate quality’ therefore varies semi-continuously with respect to the number of **D/E** residues in the target site, which makes this kinase particularly amenable to a quantitative analysis of phosphorylation stoichiometry and evolution. CK2 is also relevant for human health as it regulates several processes (apoptosis, cell cycle, DNA damage response, etc) and has been strongly implicated in cancer progression for many tissues (Duncan and Litchfield 2008; N. A. St-Denis and Litchfield 2009; Rabalski, Gyenis, and Litchfield 2016).

## Results

### Minimal sites are phosphorylated at reduced stoichiometry

For this analysis we focus on phosphosite ‘substrate quality’, which refers to the strength of a target sequence for its cognate kinase motif (Kang et al. 2013; C. J. Miller and Turk 2018). We investigate the relationship between kinase substrate quality and the stoichiometry, function, and evolution of phosphorylation, using the kinase CK2 as our model. CK2 is highly pleiotropic and phosphorylates D/E-containing peptides, a specificity that can be explained by the enrichment of positively charged amino acids on its substrate-binding surface **(Figure 1a)**. The D/E specificity of CK2 is especially strong at substrate positions +1 and +3 **(Figure 1b)**.

**Figure 1 -.**
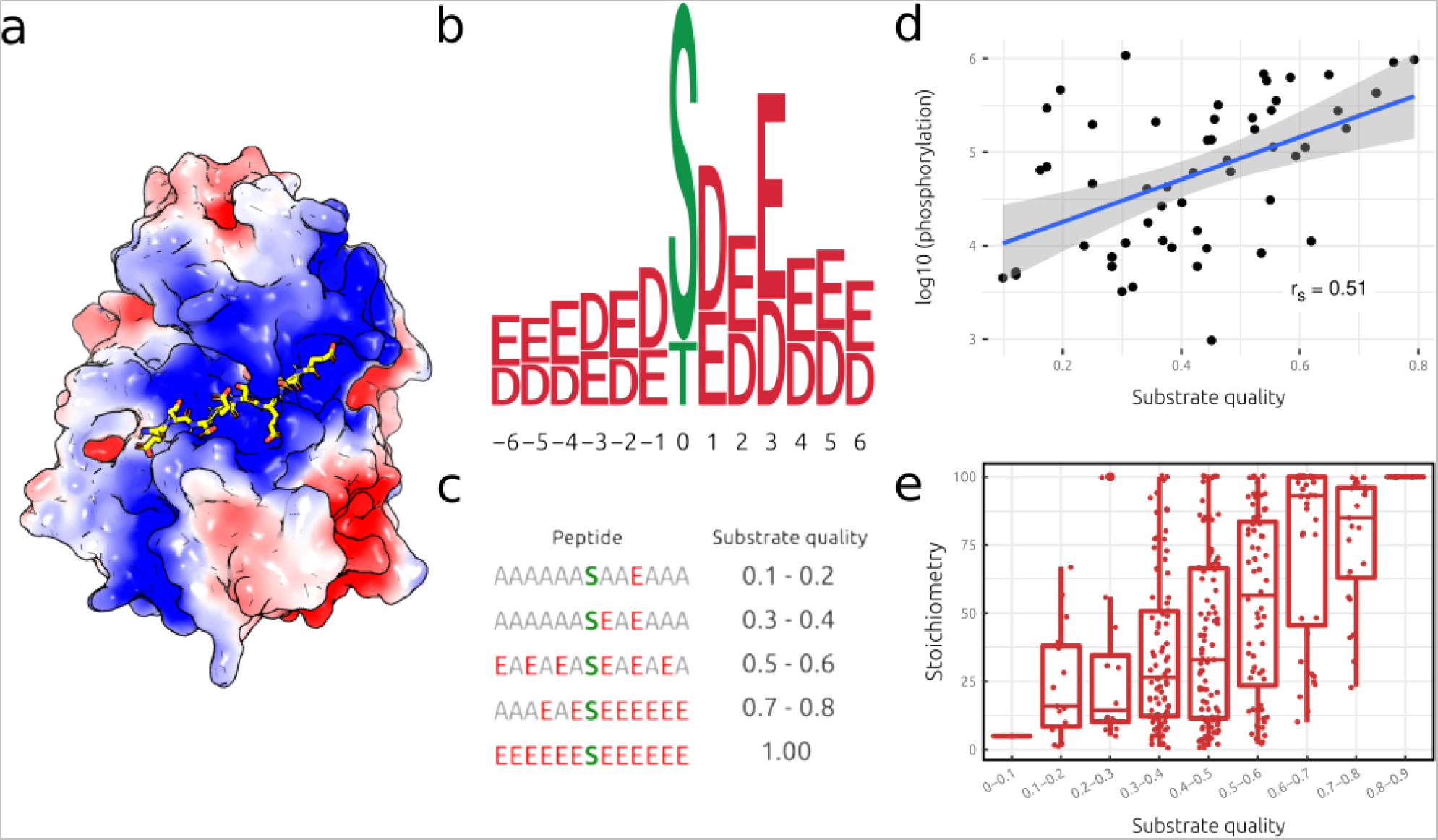
CK2 motif preference and the impact of substrate quality on the stoichiometry of phosphorylation. **a)** CK2 electrostatic surface based upon human CSNK2A1 (PDB: 3war). Peptide modelled into the substrate-binding pocket based upon DYRK1A-peptide interactions (PDB: 2wo6). Blue colour corresponds to positive charge, red to negative charge, and white to no charge **b)** Relative frequency of D and E residues among the substrates of human CK2. **c)** Weak matches to the CK2 specificity profile (b) have low substrate quality scores and vice versa. **d)** Relationship between CK2 substrate quality and phosphorylation intensity (log10 scale) for 54 WT peptides. r_s_: Spearman’s correlation. **e)** Relationship between CK2 substrate quality and the phosphorylation stoichiometry of CK2-targeted phosphosites measured in (Tsai et al. 2015).

CK2 has 581 substrates in human that have been recorded in kinase-substrate databases **(Supplementary table 1)**. This number reduces to 467 after excluding candidate multi-phosphorylation sites with a serine at positions +1 and/or +3, which can be hierarchically phosphorylated by CK2 (N. St-Denis et al. 2015). The remaining sites are enriched for D/E residues at all flanking positions **(Supplementary figure 1a**, **Figure 1b)**. To measure CK2 substrate quality, we calculate for each substrate a summed score reflecting relative D/E preferences at each flanking position (−6 to +6). For example, an ideal substrate is assigned a score of 1, an intermediate substrate a score of 0.5, and a fully minimal substrate a score of 0 **(Figure 1c).**

We start by using a peptide array assay to confirm that the intensity of phosphorylation by CK2 increases with the peptide’s substrate quality. For this purpose we selected 76 WT peptides **(Figure 1d)** that span a broad range of substrate qualities and were previously identified as *in vitro* targets for CK2 from (Tsai et al. 2015). In addition, we designed 203 mutants **(Supplementary figure 2)** to systematically increase and decrease the quality of the substrate with respect to the CK2 motif. This analysis confirms the expectation that full motif matches lead to more efficient phosphorylation and *vice versa,* for both the WT **(Spearman’s correlation = 0.51, p=8.5×10^−5^, log10-transformed)** and WT and mutant sets **(Spearman’s correlation = 0.60, p=7.8×10^−21^, log10-transformed)**. Notably, use of more complicated specificity models that account for all 20 amino acids (as opposed to just **D/E** as in **Figure 1c**) does not improve this correlation, either using known substrates to derive the specificity model **(Supplementary figure 3a)** or *in vitro* specificity data for CK2 **(Supplementary figure 3b,c)**. All peptide sequences and scores are provided in **Supplementary table 2.**

Phosphorylation stoichiometry *in vivo* is determined by the balance of kinase and phosphatase activities. We therefore questioned whether increases in substrate quality would also amount to significant changes in phosphorylation occupancy. The stoichiometry of phosphorylation for sites targeted by CK2 was measured recently on a proteome-wide scale (Tsai et al. 2015). Using this data, we indeed find a positive relationship **(Spearman’s correlation = 0.44, p=5.5×10^−18^)** between the CK2 substrate quality and the stoichiometry of phosphorylation measured in human cells **(Figure 1e).** This conclusion holds when modelling CK2 specificity using peptide array data instead of a sequence profile of known substrates (Hutti et al. 2004) **(Supplementary figure 1b)**. The CK2 substrate quality also correlates positively with kinome-wide stoichiometry for predicted CK2 substrates from three other stoichiometry datasets (Olsen et al. 2010; Wu et al. 2011; Sharma et al. 2014) **(Supplementary figure 1c)**. For the human stoichiometry data **(Supplementary figure 1c, left panel, right panel)**, we observe a weaker association between substrate quality and stoichiometry, possibly because some predicted CK2 substrates are phosphorylated by other acidophilic kinases such as the PLK and GRK families (Johnson et al. 2023). Finally, we find that CK2-specific stoichiometry is still increased by substrate quality after controlling for the negative charge on the substrate i.e. when comparing peptides with the same number of D/Es but with different substrate qualities **(Supplementary figure 4).**

To test the generality of these findings, we repeated the analysis in part for the other two kinases with available kinase-specific stoichiometry data: MAPK (Erk2) and EGFR (Tsai et al. 2015). For MAPK, the average stoichiometry is greater for optimal target sites (P-x-S/T-P) compared to minimal target sites (S/T-P) **(Supplementary figure 1d)**. However, we find no such difference between minimal and non-minimal target sites of the tyrosine kinase EGFR **(Supplementary figure 1e)**. This may reflect the stronger reliance of tyrosine kinases on accessory domains such as SH2 and SH3 domain for binding and specificity (Ubersax and Ferrell 2007).

### Several biological factors determine stoichiometry

We next constructed a generalised linear model (GLM) to quantify the relationship between CK2 stoichiometry and substrate quality. The statistical significance of substrate quality as an explanatory variable was calculated by comparing the statistical deviance between a simple model (with substrate quality as the sole explanatory variable) and a null model (without any explanatory variables). As expected, substrate quality is a highly significant explanatory variable **(p = 6.8 × 10^−22^, F-test, Figure 1e)**.

We then tested other biological features that could impact upon CK2 stoichiometry. Protein abundance has been shown previously to correlate negatively with phosphorylation stoichiometry in the yeast phosphoproteome (Levy, Michnick, and Landry 2012), and this trend is recapitulated for the CK2-specific stoichiometry data in human **(p = 4. × 10^−11^, F-test, Supplementary figure 5a)**. Predicted protein disorder also correlates positively with CK2 stoichiometry **(p = 2.9 × 10^−22^, F-test, Supplementary figure 5b)**, as shown previously in yeast for phosphoproteome-wide data (Wu et al. 2011). We also find subcellular localisation to be a correlate of CK2 stoichiometry (Wu et al. 2011), with nuclear substrates having the highest average stoichiometry **(p = 1.0 × 10^−03^, F-test, Supplementary figure 5c)**. Interestingly, while a physical interaction between the kinase and substrate is required for phosphorylation, we find a significantly higher stoichiometry if the kinase and substrate had previously been reported to interact in a database of protein-protein interactions **(p = 8.7 × 10^−04^, F-test, Supplementary figure 5d).** Finally, serines are phosphorylated at significantly higher stoichiometries than threonines for CK2 substrates **(p = 7.2 × 10^−07^, F-test, Supplementary figure 5e).**

The GLM framework allows statistical interactions between explanatory variables to be tested for. For example, an interaction was found between substrate quality and protein abundance, with high abundance substrates of low substrate quality having especially low stoichiometries **(p = 7.7 × 10^−04^, F-test, Supplementary figure 5f).** Reported physical interactions and substrate quality also combine non-additively **(p = 3.8 × 10^−02^, F-test, Supplementary figure 5g).** Finally, we observe increased stoichiometry for serine over threonine in disordered sites but not ordered sites, although this interaction was not found to be statistically significant (**p = 5.6 × 10^−02^, F-test, Supplementary figure 5h)**. Corresponding p-values for the MAPK and EGFR stoichiometry data are given in **Supplementary figure 5i-k** alongside the p-values for the CK2 stoichiometry data. All relevant data for these models are provided in **Supplementary table 3**.

In summary, this analysis demonstrates that several factors in addition to substrate quality influence stoichiometry. Other factors, such as cell cycle stage and phosphatase activity, are also very likely to be important but we lack the data to model their effects (see *Discussion*). The results also imply that the dynamics of phosphosite evolution will be impacted by the abundance of the whole protein substrate **(Supplementary figure 5a**, **Supplementary figure 5f)**.

### CK2 minimal sites are functionally distinct from optimal sites

We next determined if variation in substrate quality between phosphosites could be linked to differences in phosphosite function. We first tested the relationship between CK2 substrate quality and phosphosite functional scores (between 0 and 1) obtained from a computational predictor (Ochoa et al. 2019). To prevent circularity, the predictive model was retrained after excluding features associated with kinase specificity (see *Methods*). We find that high quality substrates tend to have higher functional scores **(Figure 2a)**, and that this trend is repeated when predicting CK2 substrates using the CK2 motif and known transient interactors of CK2 (Niinae et al. 2021) **(Supplementary figure 6a)**. For the known CK2 targets, there are a small number of weak substrates **(n=17)** with high functional scores **(Figure 2a).** These may correspond to promiscuous substrates that are functionally coupled to many kinases, but we do not find strong evidence for this when using known (albeit incomplete (Invergo and Beltrao 2018)) kinase-substrate relationships as a benchmark **(Supplementary figure 6b).**

**Figure 2 -.**
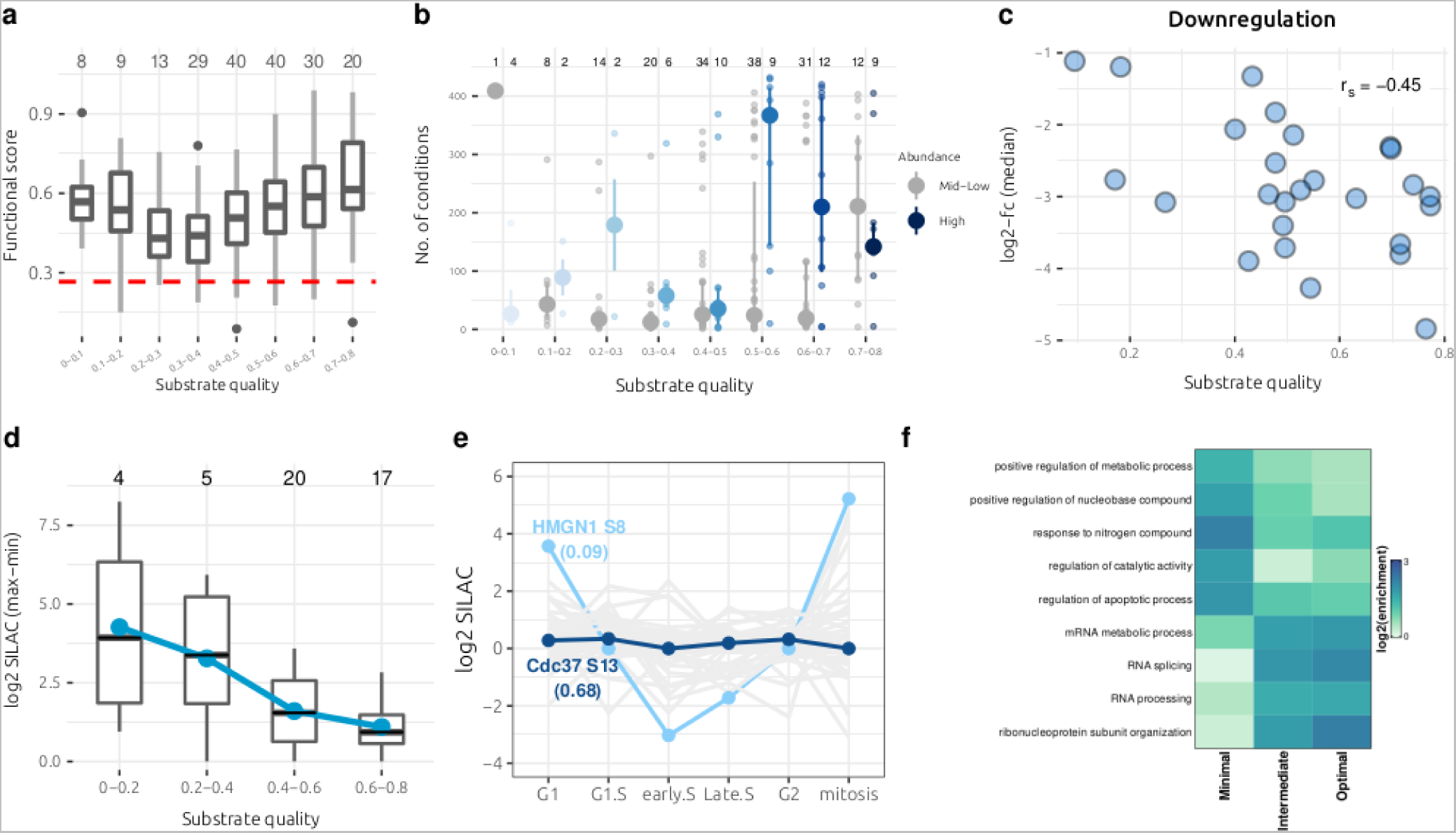
Examining the relationship between CK2 substrate quality and phosphosite function. **a)** Optimal sites have a higher functional phosphosite score than minimal substrates. Dashed red line corresponds to the median score across the human phosphoproteome. **b)** Optimal phosphosites tend to be found in more biological conditions than minimal substrates. Large dots and bars correspond to the median and interquartile range, respectively. Mid-low abundance: <100 parts per million (ppm). High abundance: >100 ppm. Dataset includes 435 conditions from (Ochoa et al. 2016). **c)** For known and predicted CK2 target sites in yeast that are downregulated in test conditions, optimal sites exhibit a stronger log2-fold decrease in phosphorylation relative to the control. Log2-fc: log2 fold-change **d)** Optimal CK2 sites tend to be less variable during the human cell cycle than minimal sites, as measured by the difference between the maximum and minimum log2 SILAC ratios across the six cell cycle stages assayed in (Olsen et al. 2010). SILAC ratios for each cell cycle stage are with respect to a reference population of asynchronous cells (Olsen et al. 2010). Blue dots correspond to the mean within each group. **e)** Phosphorylation profile across six cell cycle stages of a minimal (HMGN1 S8) and optimal (CDC37 S13) CK2 target site, in terms of log2 SILAC ratios for each stage. Log2 SILAC ratios for each cell cycle stage are with respect to a reference population of asynchronous cells (Olsen et al. 2010). The substrate qualities (SQ) for each site are in parentheses. Grey lines correspond to the phosphorylation profile of all other human CK2 substrates with SILAC measurements for each of the six cell cycle stages. **f)** Gene ontology (GO) term enrichment analysis of minimal (SQ < 0.3), intermediate (0.3 < SQ < 0.6), and optimal (SQ > 0.6) CK2 sites for biological process (BP) terms. A sample of differentially enriched terms are shown in this panel. SQ: substrate quality.

We then tested if strong CK2 substrates tend to be phosphorylated in more biological conditions, assuming that sites with higher predicted scores **(Figure 2a)** are more likely to be constitutively functional (Ochoa et al. 2019). To test this hypothesis, we analysed a global phosphoproteome dataset derived from 41 different publications and 435 conditions in human (Ochoa et al. 2016). Indeed, we find that sites with higher substrate qualities tend to be phosphorylated across a larger range of biological conditions **(Figure 2b)**. However, this may simply reflect the bias of MS/MS-based phosphopeptide detection towards more abundant phosphosites (Ochoa et al. 2019). As expected, when analysing sites **(n=2,178)** across the human phosphoproteome of known abundance and phosphorylation stoichiometry (Sharma et al. 2014), the number of detected conditions gradually increases with increasing phosphosite abundance **(Supplementary figure 7a)**. We compare this standard (i.e. proteome-wide sites of known abundance and stoichiometry) with the number of observed conditions for CK2 substrates, using the known substrate abundance and approximating stoichiometry from the substrate quality **(Figure 1e).** By comparing CK2 target sites with proteome-wide sites of similar phosphorylated abundance, we find that strong CK2 substrates are found in significantly more conditions than expected **(p=6.5×10^−4^, Mann-Whitney test, one-sided, Supplementary figure 7b, right facet)**, whereas weak substrates are found in a comparable number of conditions to sites of similar phosphorylation abundance (**p=1.8×10^−1^, Mann-Whitney test, one-sided, Supplementary figure 7b, left facet)**. We then used a recent machine learning model to predict if differences in substrate quality (SQ) affect the physical properties of phosphopeptides in terms of their ion mobility during mass spectrometry (Zeng et al. 2022), which could also explain the trend observed in **Figure 2b**. While there is a significant relationship between the two across the full range of substrate qualities **(Spearman’s correlation = 0.25, p=1.3×10^−14^, Supplementary figure 7c)**, this correlation disappears when focusing on intermediate-strong sites (SQ > 0.3; **Spearman’s correlation = 0.046, p=2.0×10^−1^, Supplementary figure 7d)** and so cannot explain the sharp increase in detected conditions observed for strong sites only (SQ > 0.6). Taken together, these data suggest that high quality CK2 sites are phosphorylated in many conditions and that technical MS/MS biases are unlikely to explain this phenomenon.

We also consider the impact of substrate quality on phosphosite regulation. CK2 is considered to be a constitutively active kinase but paradoxically has been implicated in the regulation of several signalling pathways such as the PI3K/AKT pathway, Wnt signalling pathway, and the DNA damage response (DDR) (Guerra 2006; Rabalski, Gyenis, and Litchfield 2016; Franchin et al. 2017; Borgo et al. 2021), which are activated in specific conditions. While the kinase is constitutively active, the level of kinase activity can be affected by protein interactions, CK2 quaternary structure, and PTMs via poorly understood mechanisms (Borgo et al. 2021; Roffey and Litchfield 2021). We test the relationship between CK2 substrate quality and phosphoregulation using a comprehensive map of phosphosite regulation in *S. cerevisiae* across ~100 conditions (Leutert et al. 2022). Crucially, phosphosite changes are measured on a 5-minute time scale and so likely reflect shifts in phosphorylation stoichiometry instead of protein abundance (J. Li et al. 2019; Leutert et al. 2022). We observe that, while the upregulation of CK2 substrates occurs occasionally, downregulation is much more common on this short time scale **(Supplementary figure 8).** Also, when downregulation occurs, the fold-change between test and control conditions is greater for optimal substrates than weak substrates **(Spearman’s correlation = −0.45, p=2.2×10^−02^)**, indicating stronger phosphosite downregulation for optimal substrates **(Figure 2c)**.

We next examined phosphoproteome data across multiple cell cycle stages, given previous studies implicating differential substrate quality in cell cycle regulation (Swaffer et al. 2016; Touati et al. 2018), and that CK2 has a known role in cell cycle control (Homma and Homma 2008; St-Denis, Derksen, and Litchfield 2009; Rusin, Adamo, and Kettenbach 2017). We predict that low quality substrates are more likely to be cell cycle stage-specific given the findings above **(Figure 2b)** if we consider each stage (e.g. G1, G1/S, early-S, late-S, G2, mitosis) to be a different biological condition. Using human phosphosite data across six cell cycle stages (Olsen et al. 2010), we observe that variability of the phosphosite signal (for known CK2 substrates, **n=46**) indeed decreases across the cell cycle as substrate quality increases **(Figure 2d)**. We observe the same trend at a higher sample size **(n=142)** when using knowledge of transient CK2 interactors and the CK2 motif to predict CK2 target sites (Niinae et al. 2021) **(Supplementary figure 9a).** To a partial extent these trends may be explained by greater substrate promiscuity at low substrate qualities **(Supplementary figure 9b),** reflecting evolutionary constraint upon phosphosites targeted by multiple kinases. Two examples are given in **Figure 2e** to demonstrate divergent phosphosite behaviours during the cell cycle. The first is for a minimal CK2 phosphosite on the HMGN1 protein (HMGN1 Ser 8, PKRKVS**S**A**E**GAAK). Mitotic phosphorylation of this protein promotes its dissociation from chromatin and its export from the nucleus (Louie et al. 2000; Prymakowska-Bosak et al. 2001; Cherukuri et al. 2008), facilitating chromosome condensation during mitosis (Bustin 2001; Kugler, Deng, and Bustin 2012). The second example is for CDC37 Ser 13 (VW**D**HI**E**V**SDDEDE**TH), a co-chaperone of HSP90 that helps to fold a substantial fraction of the kinome (Mandal et al. 2007; Calderwood 2015), and that is required for CK2 activity (Bandhakavi et al. 2003). The phosphorylation of S13 by CK2 is important for CDC37 function (Shao et al. 2003), and is found phosphorylated at stable levels across the human cell cycle **(Figure 2e).**

Finally, we tested for the enrichment of functional (Biological Process) GO terms among CK2 substrates. We observe differential enrichment of GO terms between minimal and optimal substrate sets **(Figure 2f**, **Supplementary figure 10, Supplementary table 4)**. Specifically, minimal substrates tend to have regulatory functions whereas optimal substrates are involved in RNA-related functions such as RNA splicing and the assembly of ribonucleoproteins. We consider the implications of this in the *Discussion* section.

### Observed CK2 substrate quality is less than the theoretical optimum under selection

We demonstrated in the section directly above that optimal substrates are often functionally distinct from minimal substrates. This implies that the flanking number of D/Es will be optimised for the function of the phosphosite and not merely sufficient for basal phosphorylation. In other words, the phosphosite fitness function (with respect to substrate quality) will not be uniform but biassed by selection towards a specific optimum. We test this hypothesis by comparing the substrate quality distribution of known substrates against the output of an evolutionary model where the fitness function (e.g. under neutrality, stabilising selection, or directional selection) can be specified.

We start by constructing a sequential-fixation model for CK2 substrate evolution, first posed in general form by Lynch and colleagues (Lynch 2020; Lynch and Trickovic 2020; Lynch and Hagner 2015). The sequential-fixation framework models the role of mutation, drift, and selection on the fixation of sites that determine a trait of interest. Each site is considered to be biallelic, with an allele that can make a positive (+) or negative (−) contribution to the trait. The mean phenotype of the trait then corresponds to the expected number of sites with the positive allele (+) (Lynch 2020; Lynch and Trickovic 2020). In this context, we consider each amino acid in the flanking region (−6 to +6) that can either increase (D or E) or decrease (any other amino acid) the substrate quality of a CK2 phosphosite **(Figure 3a)**. The model produces as an output the expected probability of genotypic states (e.g. probability of 1 D/E, 2 D/Es, 3 D/Es, etc). The results correspond to the theoretical distribution of genotypic states (either across time for the same population or across space for lineages subject to the same population-genetic parameters) for an idealised human CK2 target, given a set of mutation rates, an effective population size, and a fitness function (see *Methods*).

**Figure 3 -.**
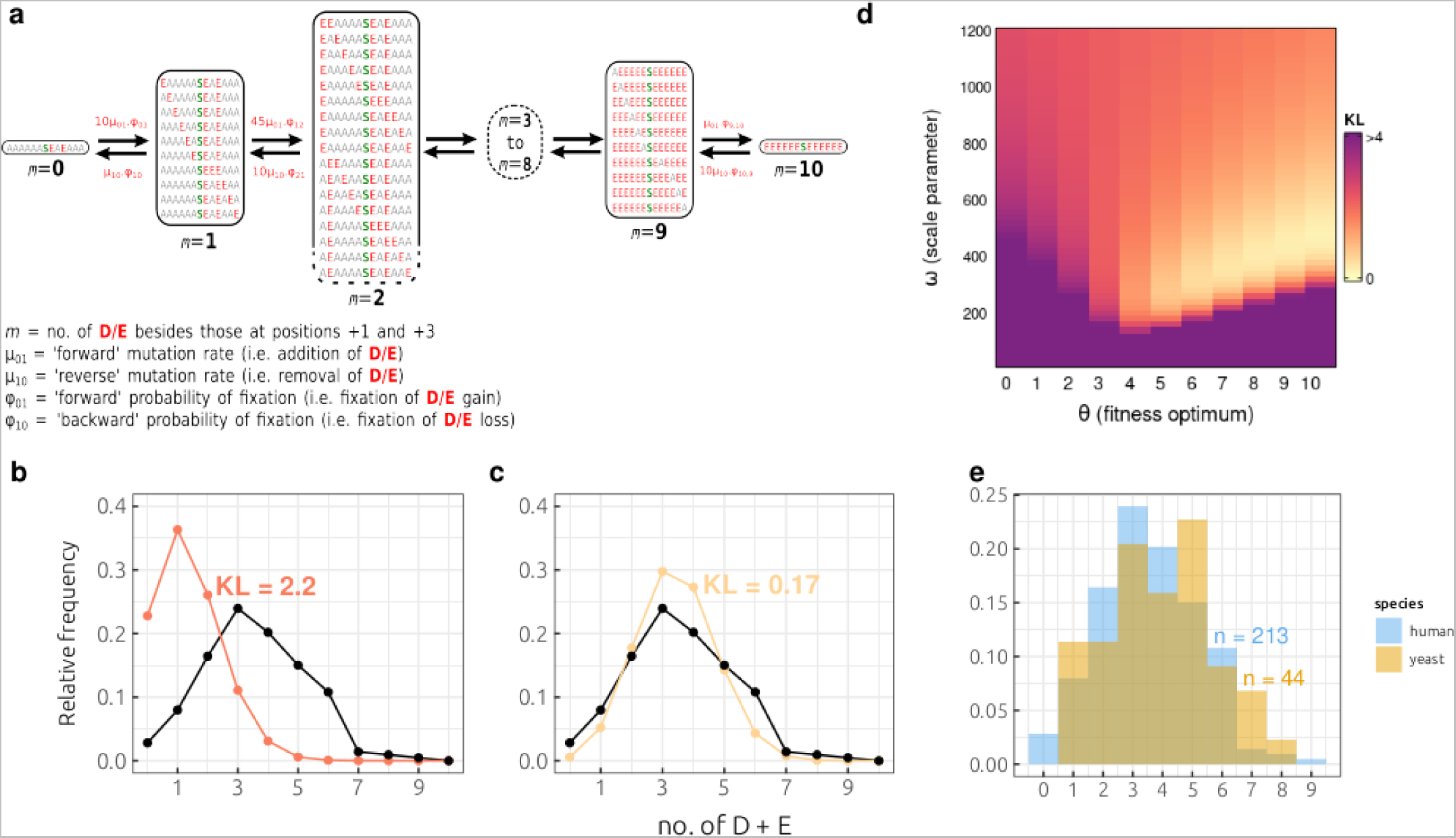
Model of CK2 target site evolution combining mutation, genetic drift, and selection. **a)** Schematic representation of a sequential-fixation model for the evolution of CK2 substrates. Model parameters are explained in the Methods section. **b)** Output of the model under selective neutrality (orange) compared to the observed distribution of D/E among known human substrates (black). KL: Kullback-Leibler divergence. **c)** Output of the model under directional selection (fitness optimum=10, scale parameter=480, yellow) compared to the observed distribution of D/E among known human substrates (black). KL: Kullback-Leibler divergence. **d)** Heatmap of the KL (Kullback-Leibler) divergence between the observed D/E distribution and that produced as an output of the sequential-fixation model. θ corresponds to the genotypic state (number of D/Es) of maximum fitness whereas ω (scale parameter) reflects the narrowness of the fitness function. The fitness function becomes narrower as ω decreases. **e)** D/E distribution for known human (blue) and yeast (yellow) substrates.

In **Figure 3b**, we present the results of the model under neutrality i.e. where a biochemically optimal site has equal fitness to a minimal site. The model assumes that positive alleles (+) from different sites will have equal effects on phosphorylation efficiency; however, this is not the case for the strongly favoured +1 and +3 positions (Hutti et al. 2004; Mok et al. 2010; Johnson et al. 2023). We therefore filter for CK2 target sites with **D/E** at +1 and +3, and then count the number of additional D/E positions in the sequence. Minimal S-D/E-X-D/E and even S-X-X-D/E peptides are sufficient for suboptimal CK2 phosphorylation *in vitro* (Marchiori et al. 1988; Sarno et al. 1997). Therefore, if selection were maintaining minimal phosphorylation status only (independent of catalytic efficiency and stoichiometry), then we would not expect the number of additional D/E residues to deviate from the expectation under neutrality. However, **Figure 3b** reveals a strong excess of D/E residues among known CK2 targets. Specifically, the observed distribution has a mean of **3.5** additional D/Es compared to an expected value of **1.4** for the theoretical value under neutrality. Moreover, **73% (155/213)** of the observed sites contain three or more additional D/Es, compared to an expectation of **15% (32/213)** under neutrality **(p = 1.0 ×10^−32^ chi-squared test)**. Overall, this suggests a significant role for natural selection in optimising phosphorylation efficiency *in vivo*.

In **Figure 3c-d** we present the results of the model under selection. To model selection we specify a Gaussian fitness function (Lynch 2020). However, the *location* (i.e fitness optimum) and *scale* (i.e relative strength of selection between genotypic states) of the fitness function are unknown **(Supplementary figure 11)**. We therefore iteratively sampled the fitness optimum (θ, 1-10 D/Es) and scale parameter (*ω*; see Methods) and then calculated the model output for all parameter combinations. In each case, the difference between the model distribution and the observed distribution was quantified using the Kullback-Leibler (KL) divergence. We find that the observed data is best explained either by a model of stabilising selection for 5-6 D/Es or directional selection for phosphosites with strong negative charges (8-10 D/Es) **(Figure 3d)**. Specifically, a fitness optimum of 6 D/Es with an *ω* value of 320 (stabilising selection) corresponds to a fitness loss of ~1.8×10^−4^ for a minimal site (S-D/E-x-S/E) relative to the optimum and gives a low KL divergence score of 0.45 **(Supplementary figure 11b)**. For directional selection, a fitness optimum of 10 D/Es and *ω* value of 480 corresponds to a fitness loss of ~2.210^−4^ for a minimal site (S-D/E-x-S/E) and also gives a low KL divergence score of 0.17 **(Figure 3c**; **Supplementary figure 11a)**. Conversely, a fitness optimum (θ) for a small number (0-3) of additional D/Es seems very unlikely given the large discrepancy between the model distribution this would produce and the empirical distribution we observe **(Figure 3d, θ = 0-3).** We also note that such a model can predict a relatively large number of minimal sites at mutation-selection equilibrium even for high fitness optima (as in **Figure 3c)**, where the number of minimal sites will depend on the strength of the mutation bias and the flatness of the fitness function.

Next, we test the prediction that the observed distribution will approach optimality as the effective population size (N_e_) increases, which will improve the efficacy of selection (Lynch and Trickovic 2020). We achieve this by comparing distributions for human (N_e_ = 2.1×10^4^) and *S. cerevisiae* (N_e_ = 7.8×10^6^). Known CK2 substrates for *S. cerevisiae* are given in **Supplementary table 5.** The two distributions overlap strongly, although we consider the analysis to be inconclusive given the small sample size of CK2 substrates (with D/E+1 and D/E+3) in yeast **(Figure 3e)**. The more complete characterisation of kinase-substrate relationships in yeast and less traditional model organisms will allow stronger inferences to be made in the future about the role of N_e_ on phosphosite evolution.

Finally, we note that the observed distribution in human has a mean of **3.5** additional D/Es whereas the sequential-fixation model predicts a likely fitness optimum of **6** or higher **(Figure 3d)**. Therefore, the distribution we observe is far from the optimal distribution we would expect if selection were operating without the constraints of genetic drift and strong mutation bias (Lynch 2018). Taken together, this computational analysis reveals a consensus view of CK2 substrate evolution where the observed substrate quality is greater than the expectation under neutrality but less than the optimum under selection.

### CK2 substrate quality is tuned by selection

Next, we compile sequence data across species to determine the extent to which minimal and optimal CK2 sites are conserved. CK2 substrate 1-to-1 orthologues were systematically retrieved, aligned, and then phylogenetically reconstructed across vertebrate species **(Figure 4a)**. We start by inferring the age of CK2 target sites, given previous studies demonstrating a relationship between phosphosite age and predicted function (Studer et al. 2016; Ochoa et al. 2019). We use ancestral sequence reconstruction across **367** CK2 sites to approximately date the emergence of a phosphorylatable residue (S or T) in the respective phylogeny of every substrate for every phosphosite position (see *Methods*). This analysis reveals that CK2 substrate phosphosites tend to be older than kinase-agnostic phosphosites phosphorylated on the same proteins **(Supplementary figure 12a)**. We next checked to see if high quality CK2 phosphosites tend to be older than low quality ones and indeed found this to be the case **(Figure 4b).** In particular, the fraction of ‘old’ sites that are at least ~600 million years old and so date to or pre-date the emergence of the vertebrates increases with substrate quality.

**Figure 4 -.**
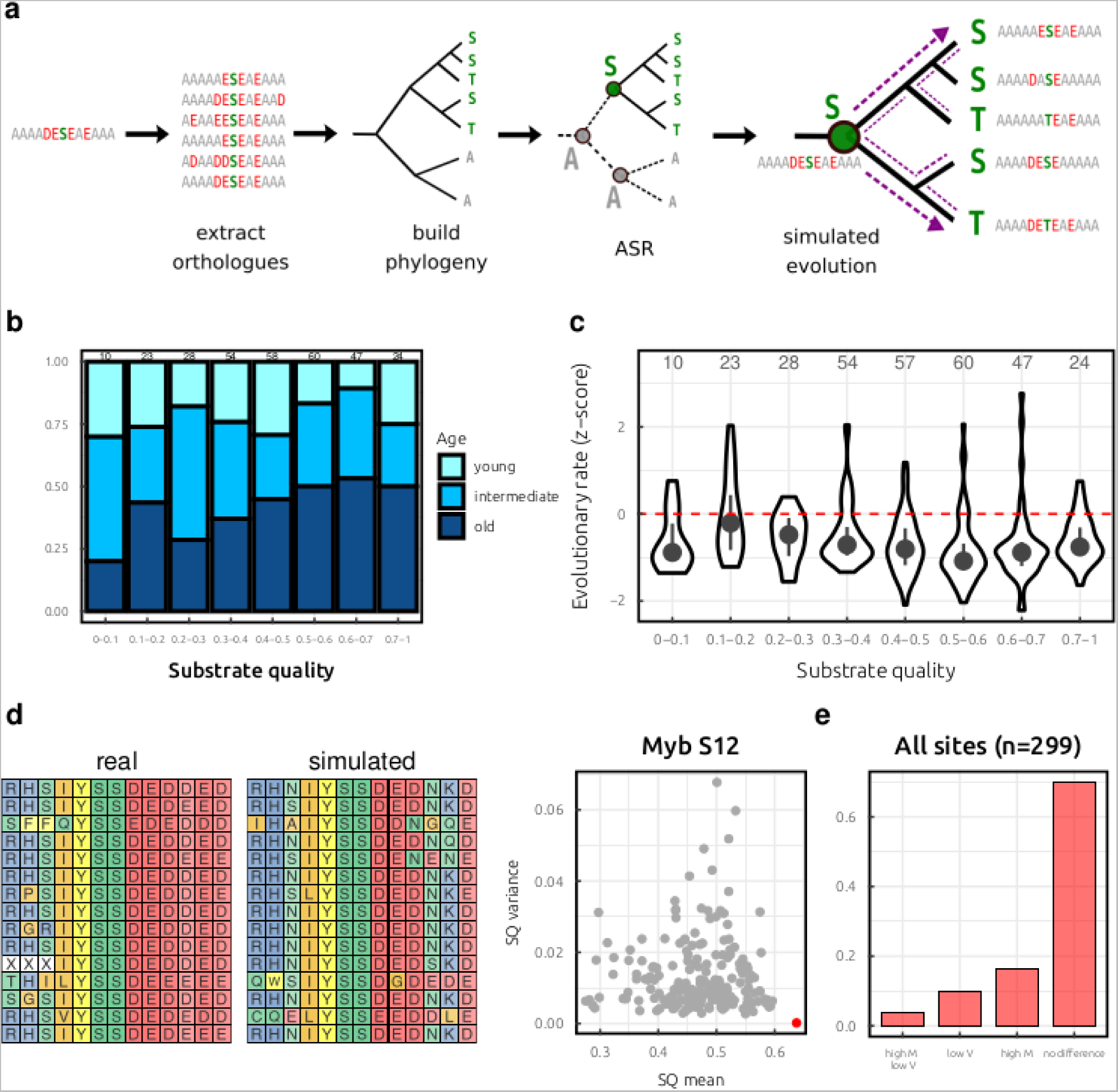
The evolution of CK2 sites across vertebrate species. **a)** For each CK2 substrate, the vertebrate one-to-one orthologues were retrieved, a phylogeny reconstructed, ancestral sequence reconstruction (ASR) performed, and simulated evolution was executed (200 trials) along the branches of the phylogeny. **b)** Estimated age of CK2 substrates with respect to substrate quality. Age was estimated conservatively from the ancestral sequence reconstructions by identifying the earliest (closest to the root) phosphorylatable residue in the phospho-acceptor position. young: <200 mya, intermediate: 200<x<600 mya, old : >600 mya. The y-axis corresponds to the relative frequency of CK2 substrates (between 0 and 1). Mya: million years ago **c)** Relationship between substrate quality and evolutionary rate. Evolutionary rate z-score normalised per protein within disordered regions. **d)** Left: multiple sequence alignment (MSA) sample for Myb S12 one-to-one orthologues. Middle: multiple sequence alignment (MSA) sample for one set of Myb S12 simulated sequences. Right: substrate quality (SQ) mean and variance for the 200 sets of simulated sequences (grey dots, n=200), and for the real set of one-to-one orthologues (red dot). **e)** Summary across 299 CK2 substrates of phosphosites with a significantly higher median (M) and lower variance (V) than the simulated sequences **(n=11)**, a significantly lower variance **(n=30)**, a significantly higher median **(n=49)**, or no significant difference detected **(n=209).** The y-axis corresponds to the relative frequency of CK2 substrates (between 0 and 1). SQ: substrate quality.

Previous studies have demonstrated that phosphosites matching a kinase recognition motif tend to be more conserved than phosphosites without any motif matches (Landry, Levy, and Michnick 2009; Beltrao et al. 2012; Freschi, Osseni, and Landry 2014). We now check for differences within a kinase motif by relating the S/T sequence conservation to the strength of the CK2 motif surrounding the phosphoacceptor. Towards this end, we calculate an evolutionary rate metric for disordered CK2 sites that normalises the rate score with respect to other disordered sites in the same substrate protein (see *Methods*). This analysis first reveals that CK2 target sites have a significantly lower evolutionary rate (i.e. are more conserved) than high-confidence phosphosites that map to the same protein but with an unknown upstream kinase **(Supplementary figure 12b)**. Importantly, we also find that the evolutionary rate decreases with increasing substrate quality **(Figure 4c),** indicating that high quality CK2 substrates are under stronger selective constraint than weak substrates. This result is consistent with the sequential-fixation model presented above predicting fitness optima at higher substrate qualities **(Figure 3d)**. CK2 targets with a substrate quality of 0-0.1 are an exception to this trend, possibly because very weak CK2 targets tend to be phosphorylated by multiple upstream kinases **(Supplementary figure 6b)**. For the results presented in **Figure 4b** and **Figure 4c**, we observe similar trends when restricting the analysis to high-confidence phosphosite alignments **(Supplementary figure 12c + Supplementary figure 12d)**, indicating that the increased age and conservation with substrate quality are not artefacts of poor quality sequence alignments (see *Methods*).

The analysis described directly above refers to the conservation of the phosphoacceptor S/T residue. We next focus on the conservation of the flanking D/E residues. While this analysis could be performed in a position-specific manner (Nguyen Ba and Moses 2010), CK2 target specificity is determined by several residues in a sequence window from positions −6 to +6 (Johnson et al. 2023) **(Figure 1b and Supplementary figure 1a).** As such, loss of substrate quality at one position may be reversed by compensatory mutations at other flanking positions. We therefore study the conservation of flanking residues by considering the substrate quality to be a quantitative trait that can be analysed using simulated evolution (Spielman and Wilke 2015; Zarin et al. 2017). Specifically, we simulate the evolution of the flanking regions from the earliest phosphorylatable (i.e. S/T-containing) ancestor to all extant descendant sequences along the branches of the phylogeny **(Figure 4a).** We then compare the substrate quality of the simulated substrates with the observed sequences extracted from the substrate orthologues **(Figure 4a).** During the simulations, we do not impose any selective constraints on the overall substrate quality but the evolution at each position is scaled by the site-specific evolutionary rates calculated from the alignments used in the **Figure 4c** analysis (see *Methods*).

Simulated evolution was performed across **299** CK2 substrates after discarding phosphosites with an insertion or deletion between the ancestral sequence (−6 to +6) and the corresponding human sequence **(Figure 4a)**. Simulated evolution was performed **200** times for each CK2 substrate, and the substrate quality mean and substrate quality variance was calculated for each replicate simulation. We then compared the obtained values with the corresponding mean and variance of the real sequences to determine if naturally evolving substrates have a significantly higher mean and/or lower variance than the neutral expectation (Zarin et al. 2019), indicating net selection above the site-specific constraints. The proto-oncogene gene Myb at S12 is an example of a CK2 target with a significantly higher mean and lower variance than the simulated sequences **(Figure 4d)**. Altogether, we find that **60** substrates have a significantly higher mean substrate quality (FDR-adjusted empirical p-value < 0.05), **41** substrates have a significantly lower variance substrate quality (FDR-adjusted empirical p-value < 0.05), with **11** substrates having simultaneously a higher mean and lower variance **(Figure 4e)**. Overall, **90/299 (30.1%)** substrates have either a significantly higher mean or lower variance, indicating evidence for selection on the CK2 substrate quality. For CK2 substrates without significant test scores, this suggests either that there is no selection on CK2 substrate quality over and above the site-specific rate constraints specified, or that there is insufficient evolutionary signal (i.e. the alignable sequences are too conserved) to detect a significant difference between the real and simulated sequences. A summary of the evolutionary results is given in **Supplementary table 6**.

Finally, we repeat the analysis above **(Figure 4d-4e)** but this time considering only secondary determinants (i.e. all sites besides **D/E+1** and **D/E+3**) on CK2 sites that also contain **D/E+1** and **D/E+3.** Since **D/E+1** and **D/E+3** are generally sufficient for phosphorylation, this approach tests for selection on determinants that improve the efficiency of phosphorylation but are not essential for it. As in **Figure 4e**, we still observe a sizable fraction of sites **(20.6%)** with substrate qualities that are maintained above the random expectation **(Supplementary figure 12e).** This indicates that such CK2 sites do not evolve simply as a binary ‘on/off’ switch but that the substrate quality is tuned by selection. Overall, our analysis of sites within **(Figure 3)** and between **(Figure 4)** species together indicate selection on the flanking sequences beyond the requirements for basal phosphorylation by CK2.

### Examination of a CK2 substrate at the kinetochore

As discussed above, the fitness function of a given site may be skewed towards intermediate or optimal motif strengths, so that the effect of any mutation will depend on the shape of the fitness function. We explore this concept further using as our case study a CK2 substrate that localises to the kinetochore during mitosis. Phosphorylation of the human +TIP CLIP-170 by CK2 increases dynactin binding, which in turn promotes the formation of stable kinetochore-microtubule attachments during M-phase (H. Li et al. 2010). The phosphosite (S1364) is of intermediate strength for the CK2 motif and found at a region of the protein predicted to be disordered (Piovesan et al. 2023) **(Figure 5a)**. For this substrate, we measured phosphorylation for 33 unique one-to-one vertebrate orthologues, 22 unique one-to-many vertebrate orthologues, and 87 systematic *CLIP-170* mutant sequences ranging from non-phosphorylatable peptides (i.e. without any D/E) to ‘ideal’ 13-mers that are saturated with negative charge (see *Methods*). Phosphorylation intensity was measured using the peptide array-based approach that was described in **Figure 1 (Figure 5b)**. This analysis reveals that phosphorylation intensity is highly conserved between one-to-one orthologues although there are a small number **(n=4)** of non-phosphorylatable orthologues. For one-to-many orthologues however the phosphorylation intensity is significantly lower on average and 41% of the peptides had no detectable phosphorylation for this assay **(Figure 5c).** This result is consistent with previous computational analyses that find widespread divergence in kinase-phosphosite relationships following gene duplication (Freschi et al. 2011; Nguyen Ba et al. 2014). For the phosphosite mutants, removing negative charge often eliminates phosphorylation completely (as expected), whereas the near-saturation of the 13-mer with negative charge leads to a 16-fold increase in phosphorylation intensity relative to the WT sequence. Therefore, the WT peptide is phosphorylated efficiently but significantly below the biophysical optimum. All peptide sequence and phosphorylation values are given in **Supplementary table 7**.

**Figure 5 -.**
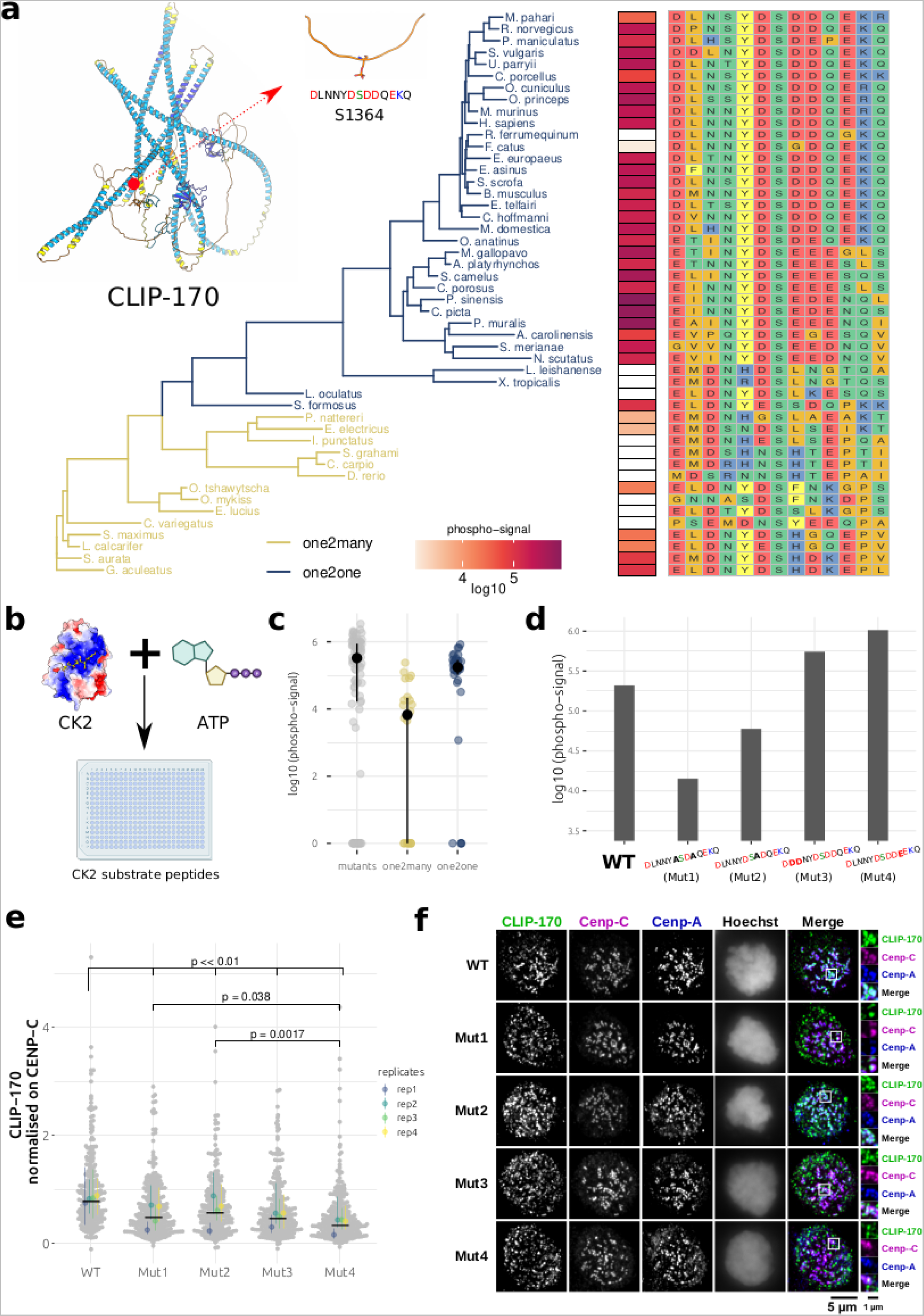
The evolution and function of a CK2 substrate at the kinetochore (CLIP-170 S1364). **a)** Sequence and phosphorylation intensity (log 10) mapped to each unique orthologue 13mer of CLIP-170 S1364. Tree visualised using ggtree and sequences visualised using ggmsa (Yu 2020; Zhou et al. 2022). **b)** Peptide array assay for quantifying peptide phosphorylation. Peptides are arrayed on a membrane, and then incubated with CK2 and radiolabeled ^32^P-ATP. The amount of phosphorylated peptide in each spot correlates with spot intensity measured using a phospho-imager. Figure panel created with the help of BioRender.com **c)** For the CLIP-170 S1364 phosphosite, difference in phosphorylation intensity (log10) for the peptide mutants, one2many orthologues, and one2one orthologues, as measured using a peptide array. Point range (black) gives the median and interquartile range. Peptides with no signal were assigned a value of 1 to give a log10-transform equal to 0.3 **d)** log10-based phosphorylation intensities for WT CLIP-170 S1364 against CK2 on a peptide array, along with four mutants designed in this study (Mut1, Mut2, Mut3, and Mut4). **e)** Fluorescence intensity for WT CLIP-170 and four mutants (Mut1, Mut2, Mut3, and Mut4). The fluorescence intensity of CLIP-170 has been normalised relative to that of the kinetochore marker protein CENP-C. The grey dots represent data points from individual kinetochores pooled across the N=4 replicates with a minimum of 15 cells measured per replicate per condition, and the coloured dots represent the median relative intensity of individual replicates. The black bar corresponds to the median (across replicates), while the point range gives the median and interquartile range (within replicates). **f)** Representative immunofluorescence images for WT CLIP-170 and four mutants (Mut1, Mut2, Mut3, and Mut4). Kinetochore markers CENP-C and CENP-A are pseudo-coloured in magenta and blue, respectively. DNA was stained with Hoechst and indicates cells in mitosis.

We next examined the effects of phosphosite mutation *in vivo*. Towards this end, we selected four mutants for further analysis: two that decrease phosphorylation below the WT value and another two that increase it **(Figure 5d)**. All mutants were tested for binding to the kinetochore by using *in vivo* fluorescence microscopy and examining CLIP-170 fluorescence relative to that of the centromere protein CENP-C, after depleting endogenous CLIP-170 with dsiRNA **(Supplementary figure 13, Supplementary table 8)**. The results quantitatively demonstrate loss of CLIP-170 localisation to the kinetochore for all four tested mutants **(p << 0.01, Tukey’s Honest test, Figure 5e and 5f)**. Moreover, the mutant that strongly increases substrate quality (Mut4) leads to a greater loss of localisation than mutants (Mut1, Mut2) that decrease it **(p = 0.038, p=0.0017, Tukey’s Honest test, Figure 5e and 5f)**. Therefore, even mutations that enhance phosphorylation *in vitro* may lead to a loss of phosphosite function in its physiological context, although the molecular mechanism for the loss of kinetochore binding is currently unknown. Both the conservation of phosphorylation *in vitro* **(Figure 5a-c)** and the in vivo mutation analysis **(Figure 5e-f)** further support our hypothesis that the CK2 flanking sequences are tuned by selection.

## Discussion

Peptide recognition modules bind peptides that have degenerate sequences that determine the extent of binding. Although sequences exist that would confer optimal binding, these are rarely observed in proteomes. Here, we examined target sites of the kinase CK2 to determine the relationship between the quality of the substrate (for the CK2 motif) and the functional properties of the phosphosite.

The main conclusions of this study can be summarised as follows: strong CK2 targets are found at higher stoichiometries, have higher functional scores, and are phosphorylated in more biological conditions than weak substrates. Conversely, weak CK2 substrates are more variable during the cell cycle and are more likely to have regulatory functions. Strong CK2 substrates tend to be older and more conserved (S/T conservation) than weak substrates and evolutionary simulations indicate that CK2 flanking sequences are often (~30% of sites) tuned by selection. The CK2 substrate quality of known human substrates is on average greater than that expected under selective neutrality but less than the theoretical optimum predicted by our evolutionary model. However, it is not currently clear whether the existence of some minimal sites can be best explained simply by mutation bias or individual fitness functions that deviate from a consensus fitness function that optimises for high substrate quality **(Figure 3)**. Our experimental characterisation of one CK2 target at the kinetochore (CLIP-170 S1364) confirmed that the phosphosite function *in vivo* can be highly sensitive to mutations at the flanking regions that increase or decrease substrate quality, a result that is consistent with our evolutionary analyses.

This study represents a kinase-centric analysis of phosphorylation dynamics that depends heavily upon the functional and bioinformatic annotations of substrates for a well characterised kinase. However, a fully quantitative understanding of phosphorylation requires knowledge of both the writer (kinase) and eraser (phosphatase) enzyme activity. Unfortunately, a more holistic approach is limited by the relative sparsity of known phosphatase-substrate relationships. To illustrate this, consider that the current release of the DEPhOsphorylataion Database (DEPOD) contains ~980 phosphatase-phosphosite annotations (Damle and Köhn 2019) compared to the ~14,400 kinase-phosphosite annotations present in the dataset used in this study. Of the ~580 known CK2 target sites in human, only 12 have been site-specifically annotated with their cognate phosphatase in DEPOD (Damle and Köhn 2019). Further advances in our understanding of phosphatase specificity (Godfrey et al. 2017; Brautigan and Shenolikar 2018; Nilsson 2019; Hoermann et al. 2020; Hoermann and Köhn 2021; Hein et al. 2023), and how this relates to phosphorylation/dephosphorylation kinetics (Godfrey et al. 2017; Touati et al. 2018; H. Nguyen and Kettenbach 2023), will hopefully allow for more integrative bioinformatic approaches in the future.

Our computational analysis of CK2 target sites suggest that low quality sites tend to be more dynamic during the cell cycle **(Figure 2d-e**, **Supplementary figure 9)** and they are also enriched for regulatory functions, as determined using GO term enrichments **(Figure 2f**; **Supplementary figure 10).** Conversely, optimal substrates are relatively static during the cell cycle but can be downregulated in some conditions **(Figure 2d-e**, **Supplementary figure 8).** When they are downregulated following a stimulus, their fold-change downregulation (test vs. control) is also greater than for weak substrates **(Figure 2c)**. One potential explanation is that strong substrates represent constitutive – possibly inhibitory – phosphosites that are downregulated under specific conditions. We consider this to be one part of a broader question of the general relationship between phosphorylation stoichiometry and function, motivated in part by previous observations that tyrosines tend to be phosphorylated at low stoichiometry, proline-directed and basic motifs at intermediate stoichiometries, and acidic motifs (like CK2) at higher stoichiometries (Wu et al. 2011; Sharma et al. 2014; Tsai et al. 2015, 2022). A previous study also found that dynamically upregulated sites during *Xenopus* egg activation have lower starting (i.e. pre-stimulus) stoichiometries than phosphosites that are unaffected by egg activation, further linking stoichiometry with function (Presler et al. 2017). Future insights could benefit from experimental mutagenesis approaches that explore the phosphosite fitness function with respect to stoichiometry and/or substrate quality.

The extent to which the insights generated here can be applied to other kinase-substrate relationships or even domain-peptide interactions in general is another important consideration. For this study we define the strength of the substrate almost exclusively in terms of the intrinsic specificity of the CK2 catalytic subunit for the region surrounding the phosphosite. However, phosphorylation specificity for other kinases is determined also by ancillary factors such as docking interactions, scaffold/adaptor interactions, and kinase PTMs (C. J. Miller and Turk 2018; Balasuriya et al. 2020; Cullati et al. 2022). For example, Dyla and colleagues recently examined how the strength of a docking interaction (i.e. away from the catalytic site) between a kinase and substrate affects the efficiency of kinase phosphorylation. Remarkably, they showed that the efficiency of phosphorylation by PKA is maximal at intermediate docking strengths and that docking is most effective when the phosphosite motif is minimal (Dyla et al. 2022). Multi-site phosphorylation is also an important factor such that overall substrate strength can reflect the combinatorial contribution of many phosphosites (and docking sites) in a narrow region of the substrate primary structure. This has been extensively characterised for the CDK family of kinases (Örd et al. 2019). Most recently, a synthetic biology approach was used to show how modulating the strength of CDK substrates – in terms of the phosphoacceptor, flanking regions, docking motifs, and inter-psite spacing – affects the dynamics of nuclear import and export (Faustova et al. 2022).

Evolution of cellular features in which non-optimal biophysical binding could be optimal for fitness could extend to many other contexts such as to other peptide-recognition modules. For example, the Sho1-Pbs2 SH3 domain interaction in yeast is maintained at relatively low affinity, and increasing the affinity of the ligand by mutation leads to a loss of fitness and increased cross-reactivity with other SH3 domains (Zarrinpar, Park, and Lim 2003). Likewise, generation of an artificial SH2 ‘superbinder’ by mutation was found to be deleterious for pTyr-based signalling (Kaneko et al. 2012). Even for protein-DNA interactions, a specificity-affinity trade-off has been posed for transcription factor binding sites (TFBS), allowing transcription factor paralogues to be discriminated by low affinity target sites (Kribelbauer et al. 2019). Selection for suboptimal binding affinity is therefore likely to be a general biological paradigm that we explore systematically here for a kinase with fundamental importance for cell biology.

## Supplementary figures

**Supplementary figure 1 -.**
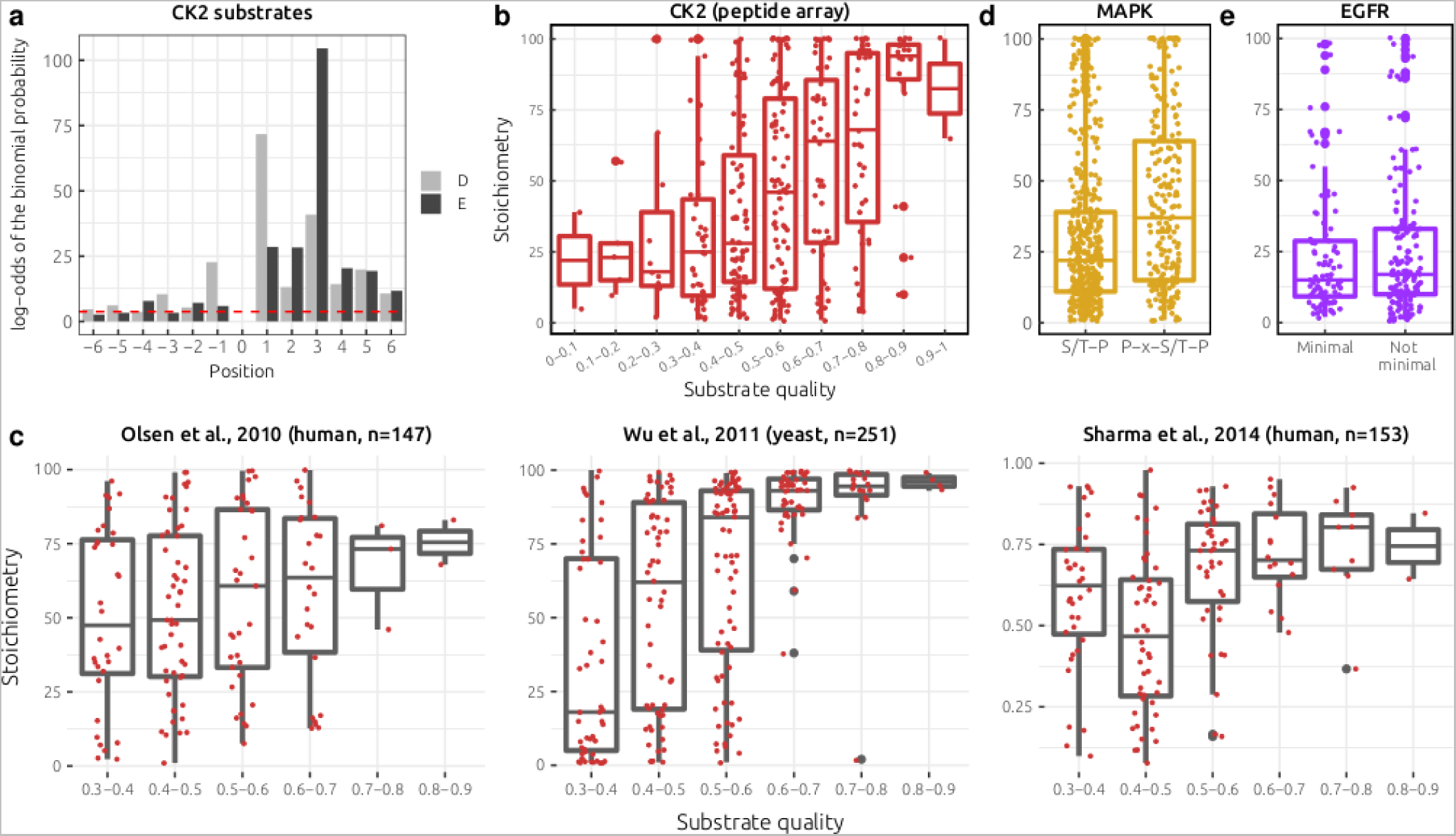
CK2 specificity and phosphorylation stoichiometry for CK2, MAPK, and EGFR targets. **a)** log-odds of the binomial probability for D and E residues being from positions −6 to +6 for known human CK2 substrates. The log-odds of the binomial probability measures the enrichment of D/E residues above what is expected given their frequency in the background proteome. Log-odds above the red line (y=3.75) correspond to a p-value < 0.05 **b)** Relationship between CK2 substrate quality and the phosphorylation stoichiometry of CK2-targeted phosphosites measured in (Tsai et al. 2015). Here the CK2 specificity was modelled using peptide array data from (Hutti et al. 2004). **c)** Relationship between CK2 substrate quality for predicted CK2 substrates and phosphorylation stoichiometry derived from (Olsen et al. 2010) (left), (Wu et al. 2011) (middle), and (Sharma et al. 2014) (right). CK2 substrates were predicted on the basis of their match to the CK2 S-D/E-x-D/E consensus. **d)** Difference in stoichiometry between ‘minimal’ MAPK sites (S/T-P) and ‘optimal’ MAPK sites (P-x-S/T-P), as measured in (Tsai et al. 2015). **e)** Difference in stoichiometry between ‘minimal’ and ‘not minimal’ EGFR sites, as measured in (Tsai et al. 2015). Distinction between EGFR minimal and ‘non minimal’ sites was made based upon the specificity model presented in (Cantor, Shah, and Kuriyan 2018).

**Supplementary figure 2 -.**
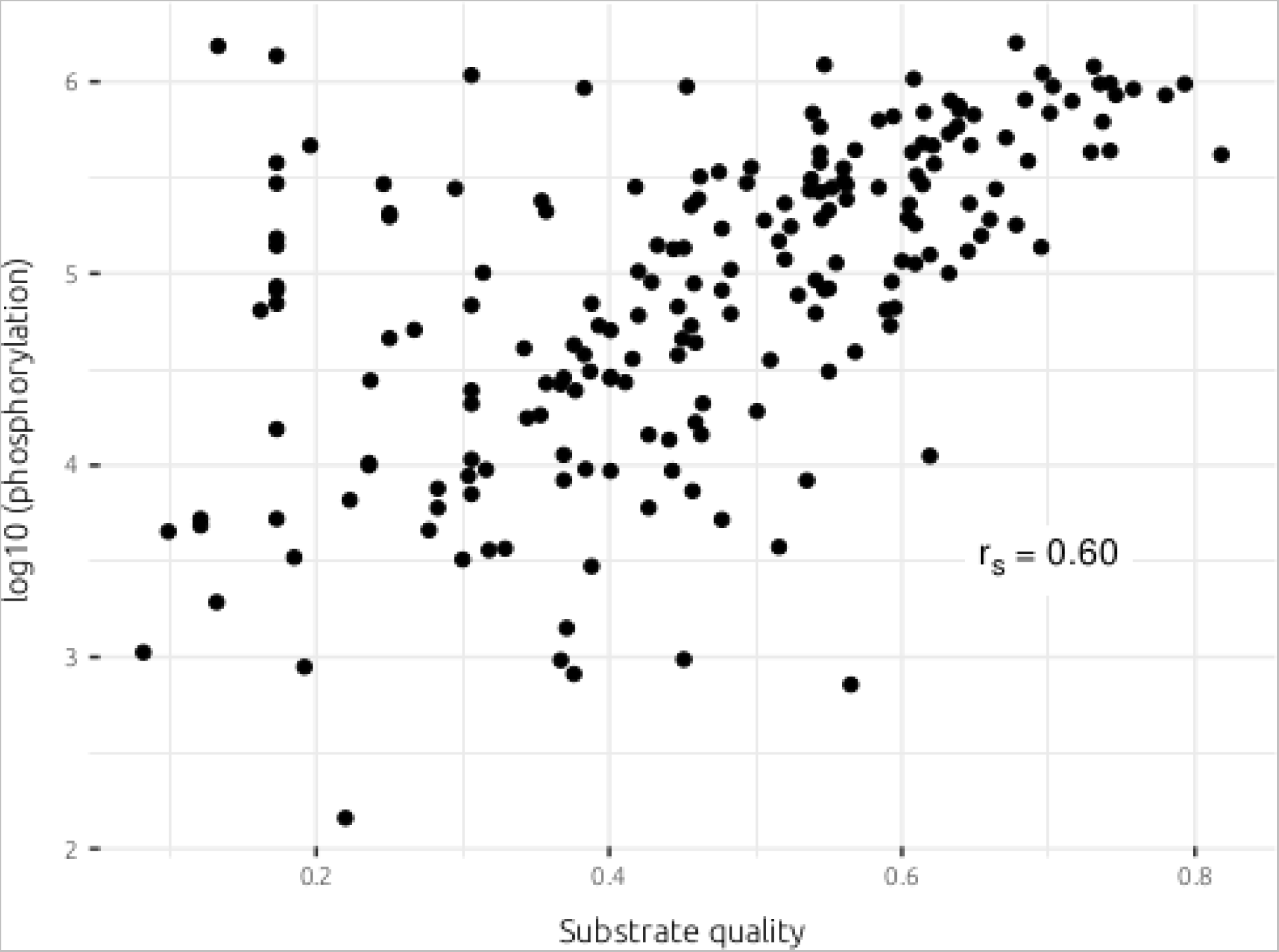
Relationship between CK2 substrate quality and phosphorylation intensity. Phosphorylation intensity (log10 scale) for 54 WT peptides and 144 mutant peptides. Phosphorylation intensity was assayed using a peptide array (see Methods). r_s_ = Spearman’s correlation.

**Supplementary figure 3 -.**
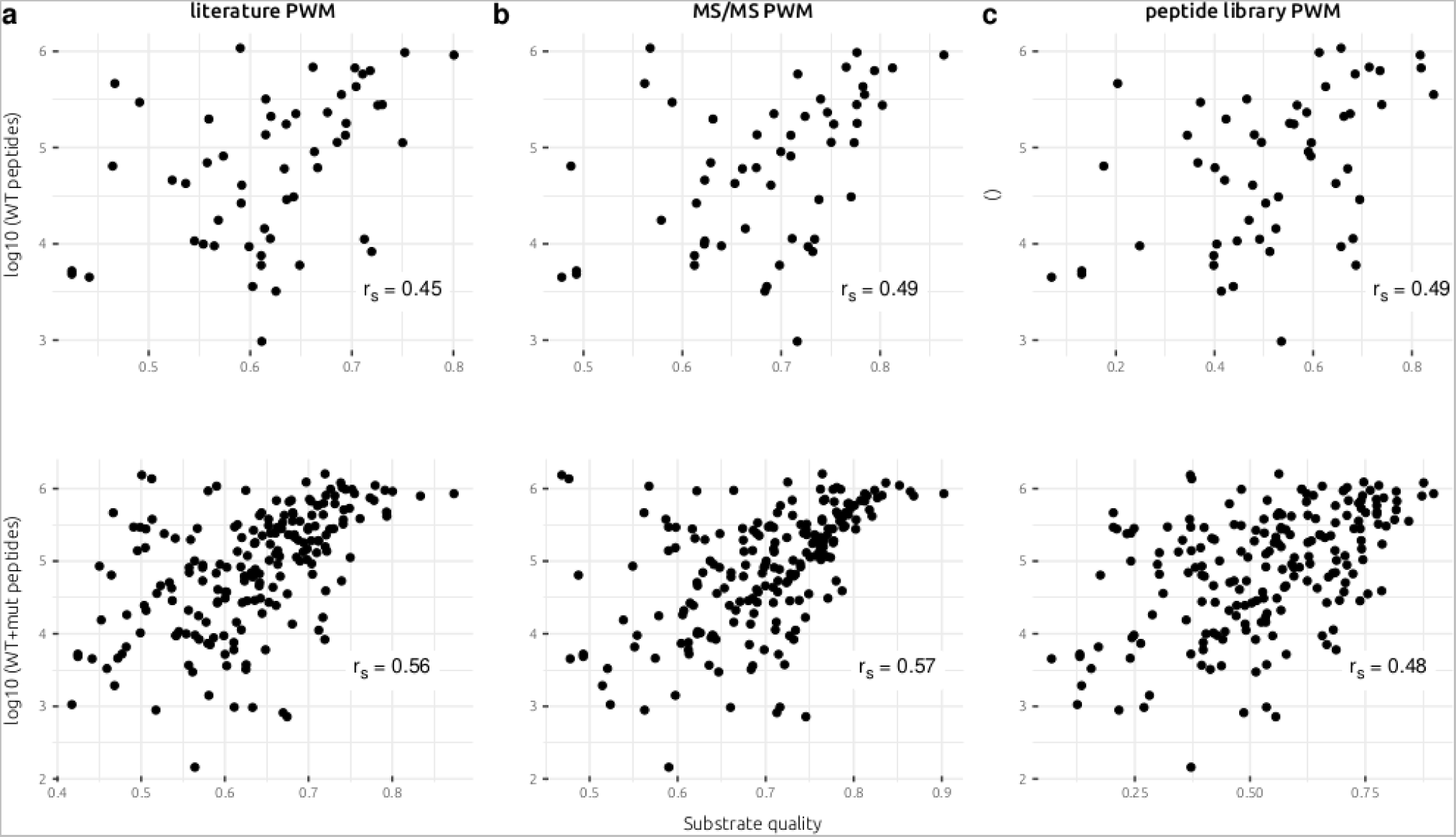
Relationship between CK2 substrate quality and phosphorylation intensity using different methods to model CK2 specificity. Phosphorylation intensity (log10) for 54 WT peptides (top row) and the combined set of 54 WT peptides and 144 mutant peptides (bottom row), all sourced from CK2-targeted peptides recorded in (Tsai et al. 2015). In panels **a**, **b**, **c** the substrate quality score is derived from different sources of data that were used to construct the position weight matrix (PWM). **a)** Known CK2 substrates recorded in (Bachman, Gyori, and Sorger 2022) were used to construct a literature-based PWM. **b)** CK2 targets phosphorylated in vitro from a proteome lysate (Sugiyama, Imamura, and Ishihama 2019) were used to construct a MS/MS-based PWM. **c)** Phosphorylation of degenerate peptide arrays in vitro was used to construct a peptide library PWM (Johnson et al. 2023). r_s_ = Spearman’s correlation.

**Supplementary figure 4 -.**
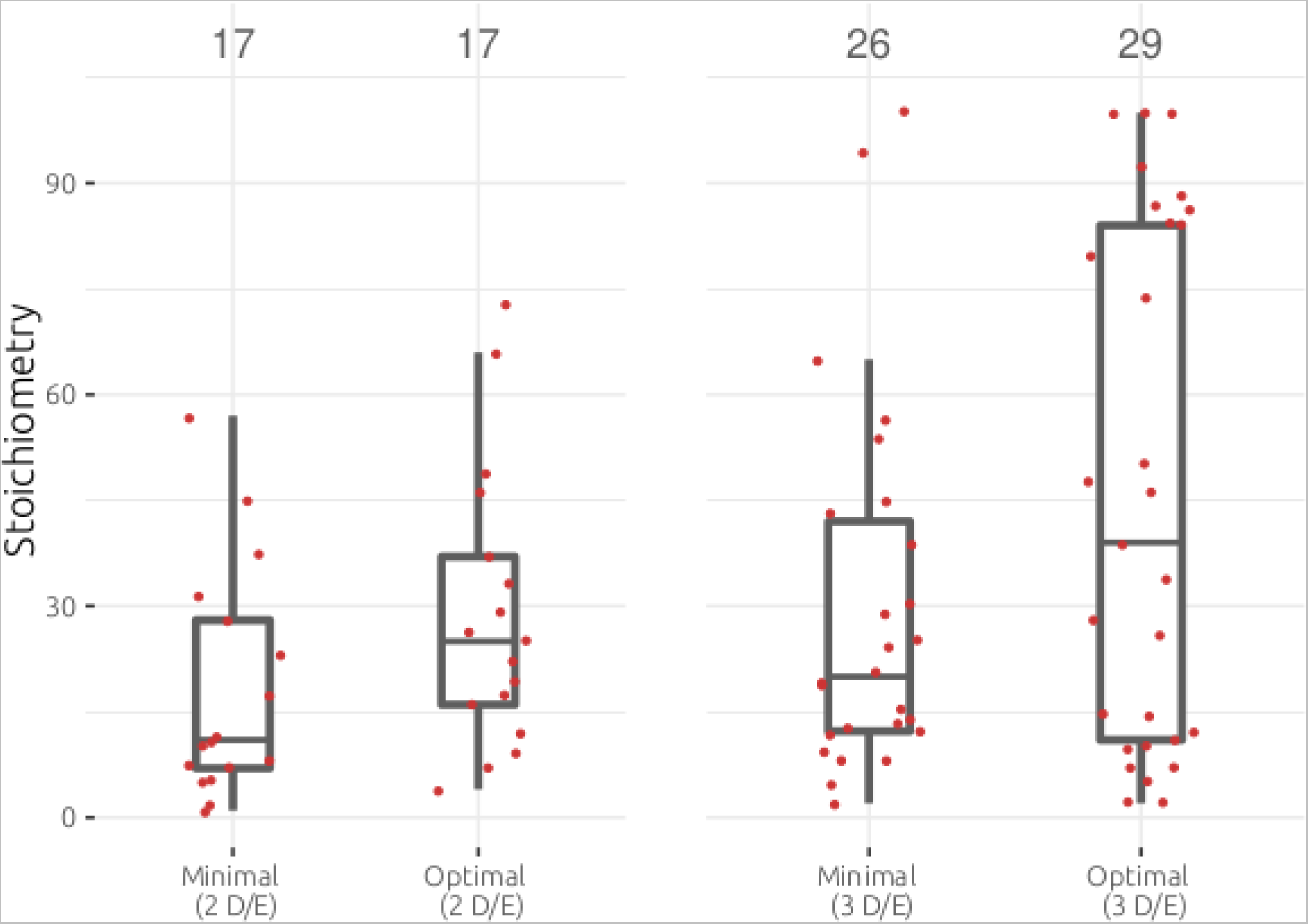
Comparing CK2 target sites with the same negative charge but different substrate qualities. For the phosphosites presented in Figure 1e, analysis of those sites with either 2 D/E residues (left) from positions −6 to +6, or 3 D/E residues (right). ‘Minimal’ here refers to sites that do not match the S-D/E-x-D/E consensus motif whereas sites labelled as ‘Optimal’ in this figure panel match the S-D/E-x-D/E consensus motif.

**Supplementary figure 5 -.**
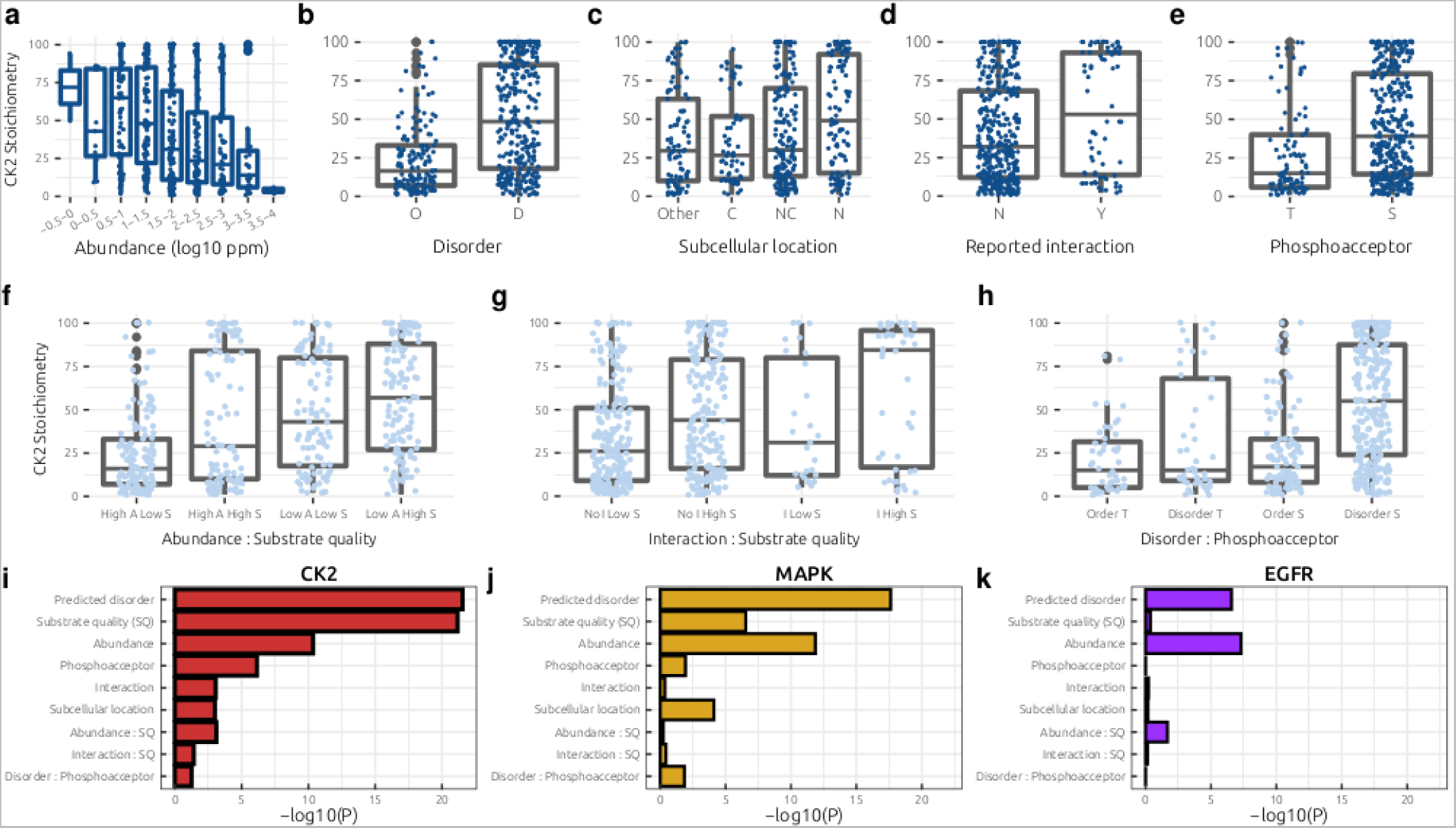
A generalised linear model (GLM) for phosphorylation stoichiometry. Association between the stoichiometry of CK2-targeted sites and **a)** substrate abundance obtained from PaxDb. Ppm: parts per million **b)** substrate order/disorder, ‘O’/’D’ **c)** substrate subcellular localisation, **C:** cytoplasm, **NC:** nucleus and cytoplasm, **N:** nucleus, **Other:** any other compartment **d)** whether the kinase and substrate have previously been reported to interact in a database of protein-protein interactions **e)** whether the phosphoacceptor is a ‘T’ or ‘S’ **f)** statistical interaction between abundance and substrate quality **g)** statistical interaction between detection of a physical interaction (between kinase and substrate) and substrate quality **h)** statistical interaction between the phosphoacceptor identity and predicted disorder **i-k)** −log10(p-values) derived from the generalised linear models (GLMs) for the **i)** CK2, **j)** MAPK, and **k)** EGFR stoichiometry data.

**Supplementary figure 6 -.**
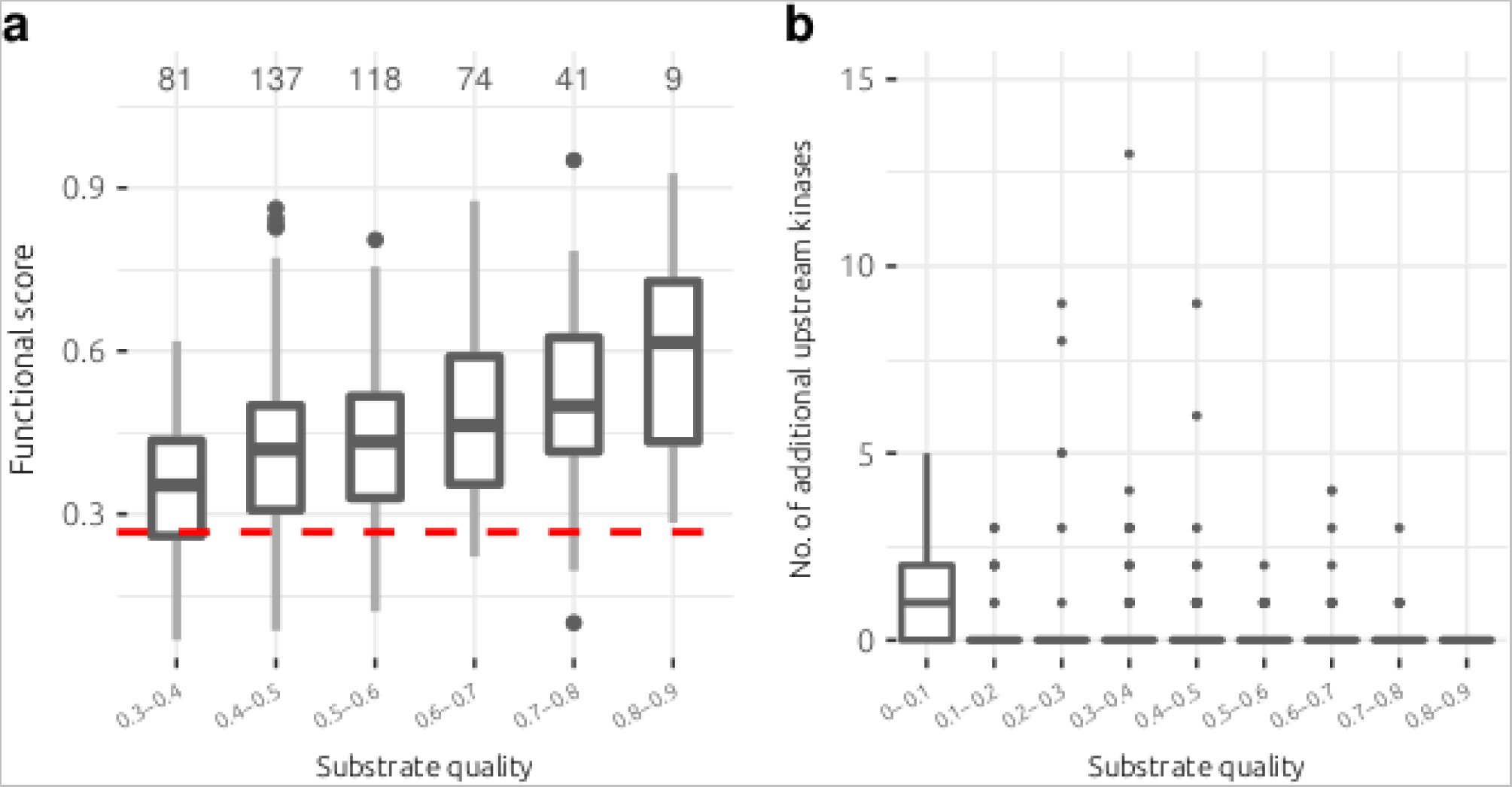
Relationship between CK2 substrate quality and functional score and substrate promiscuity. **a)** Predicted CK2 target sites with a higher substrate quality tend to have higher phosphosite functional scores. Predictions made on the basis of the CK2 substrate motif and known CK2 interactors in human. Dashed red-line corresponds to the median score across the human phosphoproteome. **b)** Relationship between CK2 substrate quality and substrate promiscuity of CK2 targets for known kinase-substrate relationships.

**Supplementary figure 7 -.**
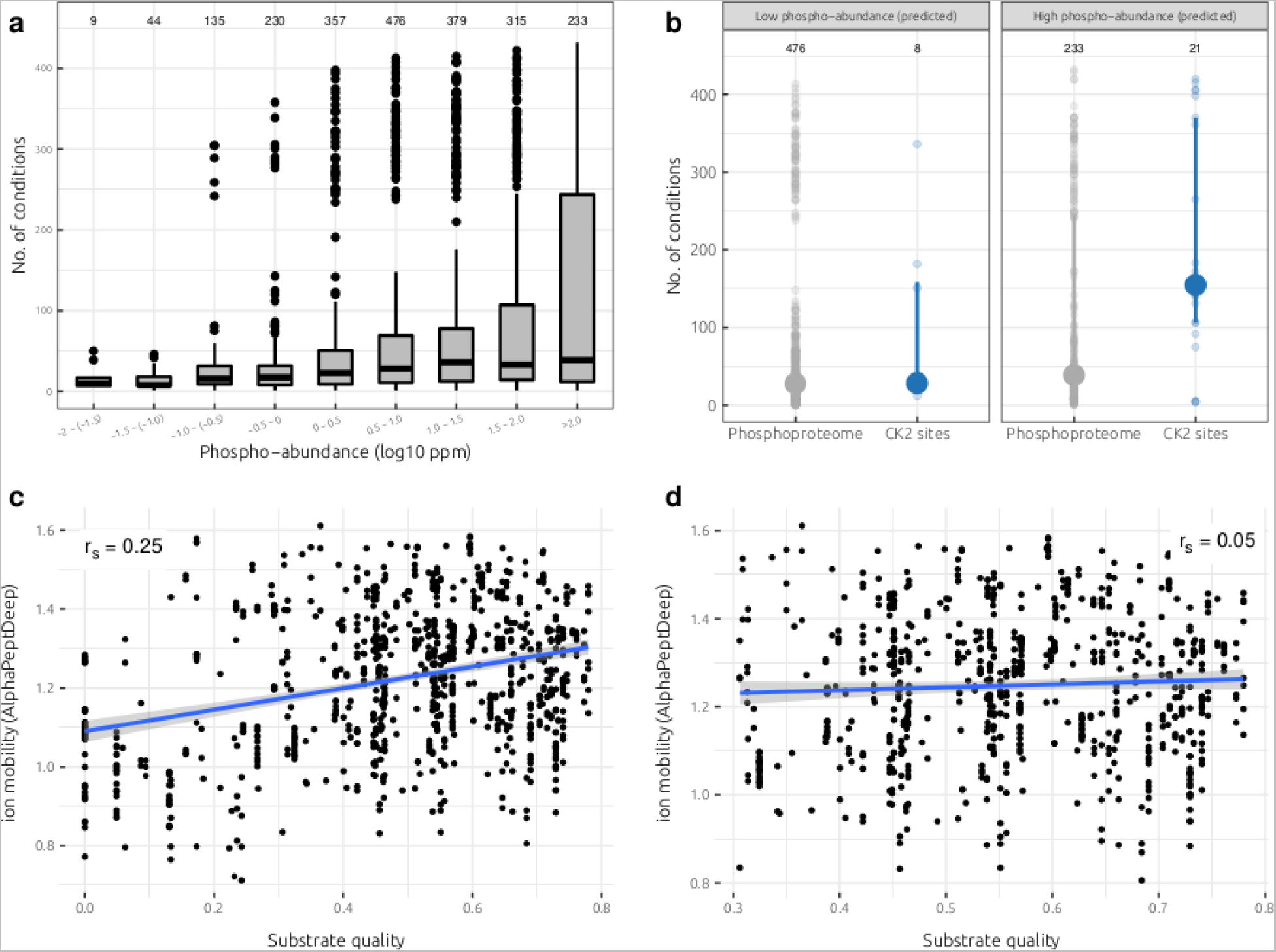
Analysing the role of phosphorylation abundance and ion mobility on the number of detected conditions. **a)** Relationship between phosphosite abundance and the number of detected conditions for phosphosites (proteome-wide) of known stoichiometry and abundance derived from (Sharma et al. 2014). Phospho-abundance = protein abundance × phosphorylation stoichiometry. Dataset includes 435 conditions from (Ochoa et al. 2016). **b)** Left grey: phosphoproteome sites of low phosphorylation abundance (3-10 ppm), left blue: CK2 target sites of low <u>predicted</u> phosphorylation abundance (abundance > 100 ppm and SQ: 0.0-0.3), right grey: phosphoproteome sites of high phosphorylation abundance (>100 ppm), right blue: CK2 target sites of high <u>predicted</u> phosphorylation abundance (abundance > 100 ppm and SQ: 0.6-0.8). **c)** Relationship between substrate quality (SQ = 0.0-1.0) of CK2 sites and ion mobility predicted for the corresponding phosphopeptides using AlphaPeptDeep (Zeng et al. 2022). r_s_ = Spearman’s correlation. **d)** Relationship between substrate quality (SQ = 0.3-1.0) of CK2 sites and ion mobility predicted for the corresponding phosphopeptides using AlphaPeptDeep (Zeng et al. 2022). r_s_ = Spearman’s correlation. SQ: substrate quality.

**Supplementary figure 8 -.**
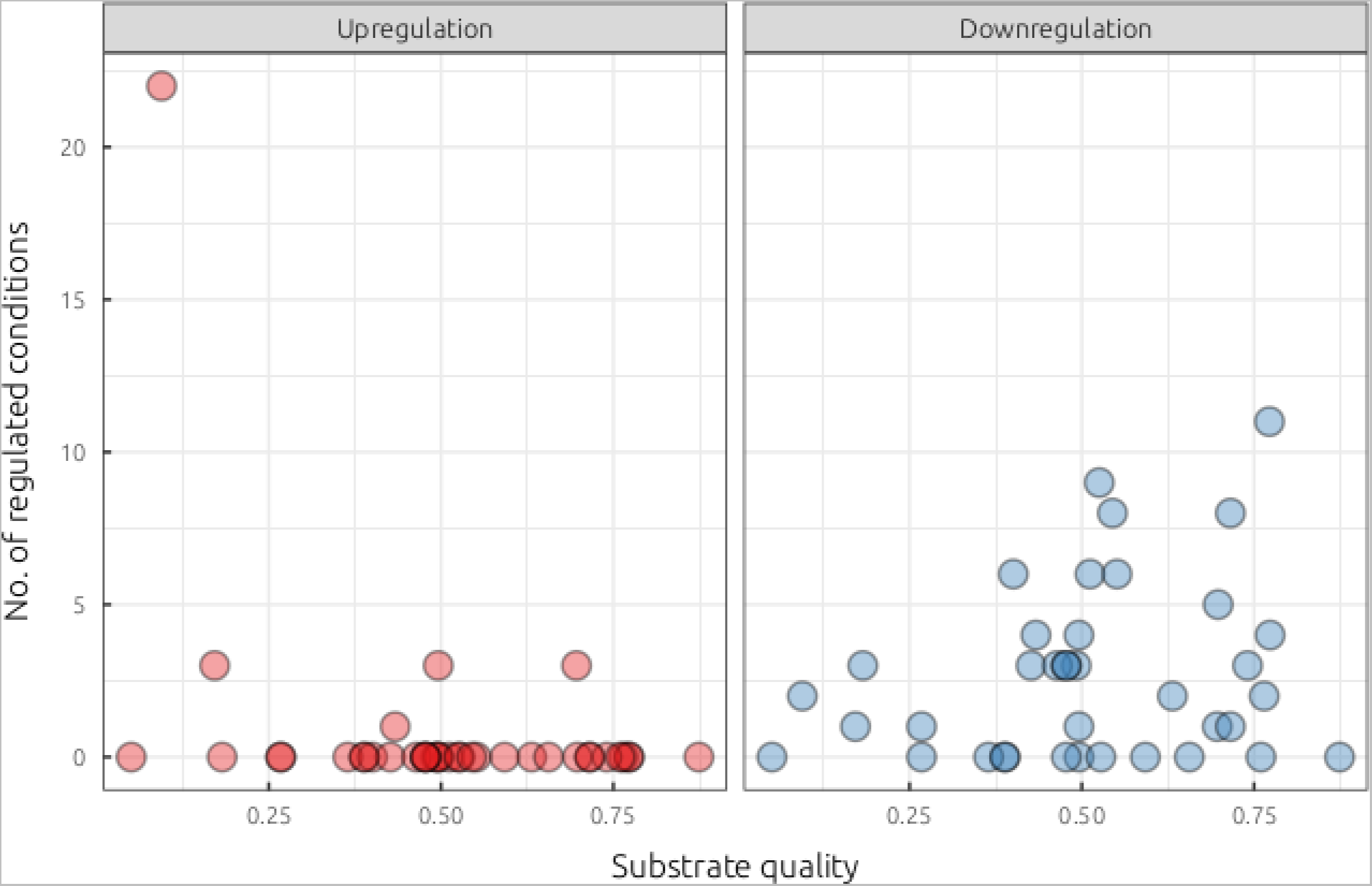
Relationship between substrate quality and the number of regulated conditions in yeast. For known and predicted CK2 substrates, the number of conditions where the phosphosite was found to be either upregulated (left) or downregulated (right) across ~100 conditions in S. cerevisiae on a 5-minute time scale, as reported in (Leutert et al. 2022). Known substrates were sourced from the literature and combined with predictions made using cka1/cka2 interactors with phosphosite matches to the S-D/E-x-D/E motif.

**Supplementary figure 9 -.**
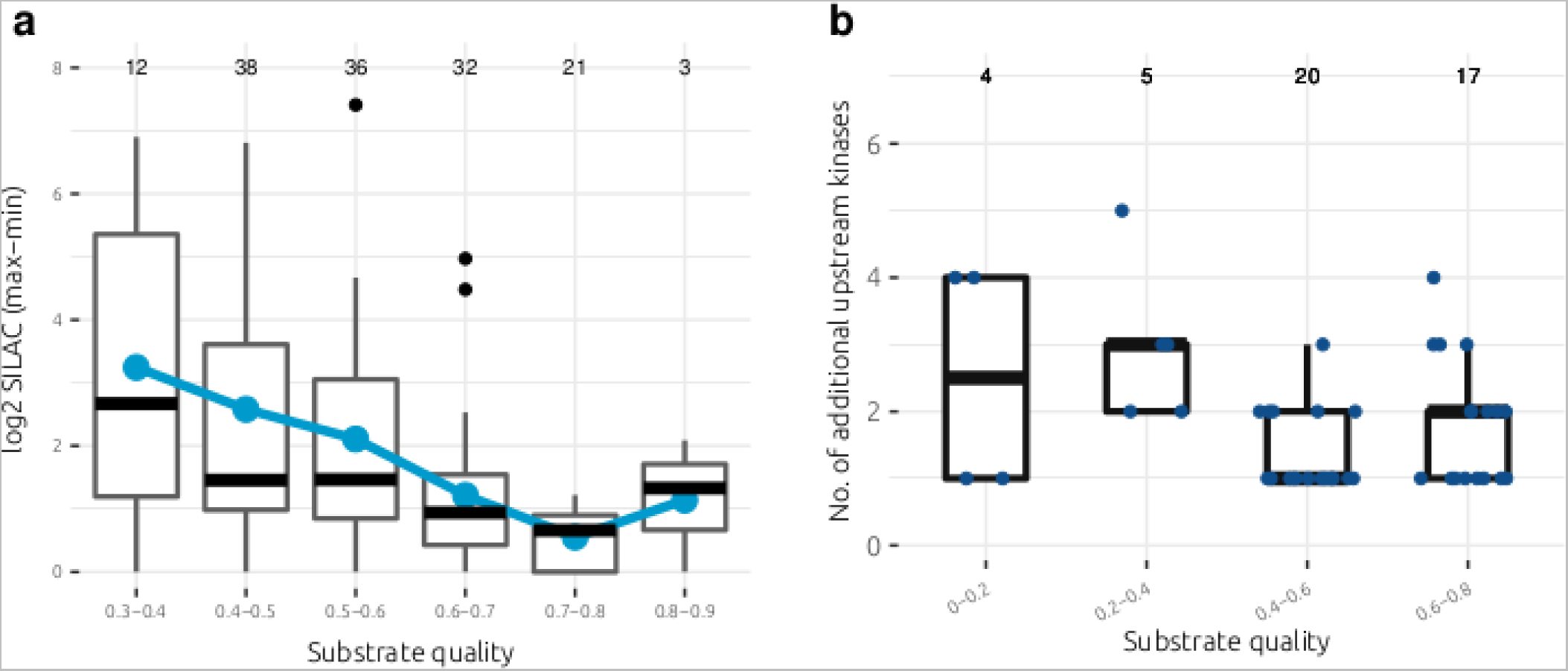
Relationship between substrate quality and cell cycle variability in human. **a)** Relationship between CK2 substrate quality and variability during the human cell cycle for predicted CK2 substrates. Predictions made on the basis of the CK2 substrate motif and known CK2 interactors in human. Cell cycle variability measured by the difference between the maximum and minimum log2 SILAC ratios across the six cell cycle stages assayed in (Olsen et al. 2010). SILAC ratios for each cell cycle stage are with respect to a reference population of asynchronous cells (Olsen et al. 2010). Blue dots correspond to the mean within each group **b**) For the known CK2 substrates visualised in Figure 2d, relationship between substrate quality and substrate promiscuity among known kinase-substrate relationships.

**Supplementary figure 10 -.**
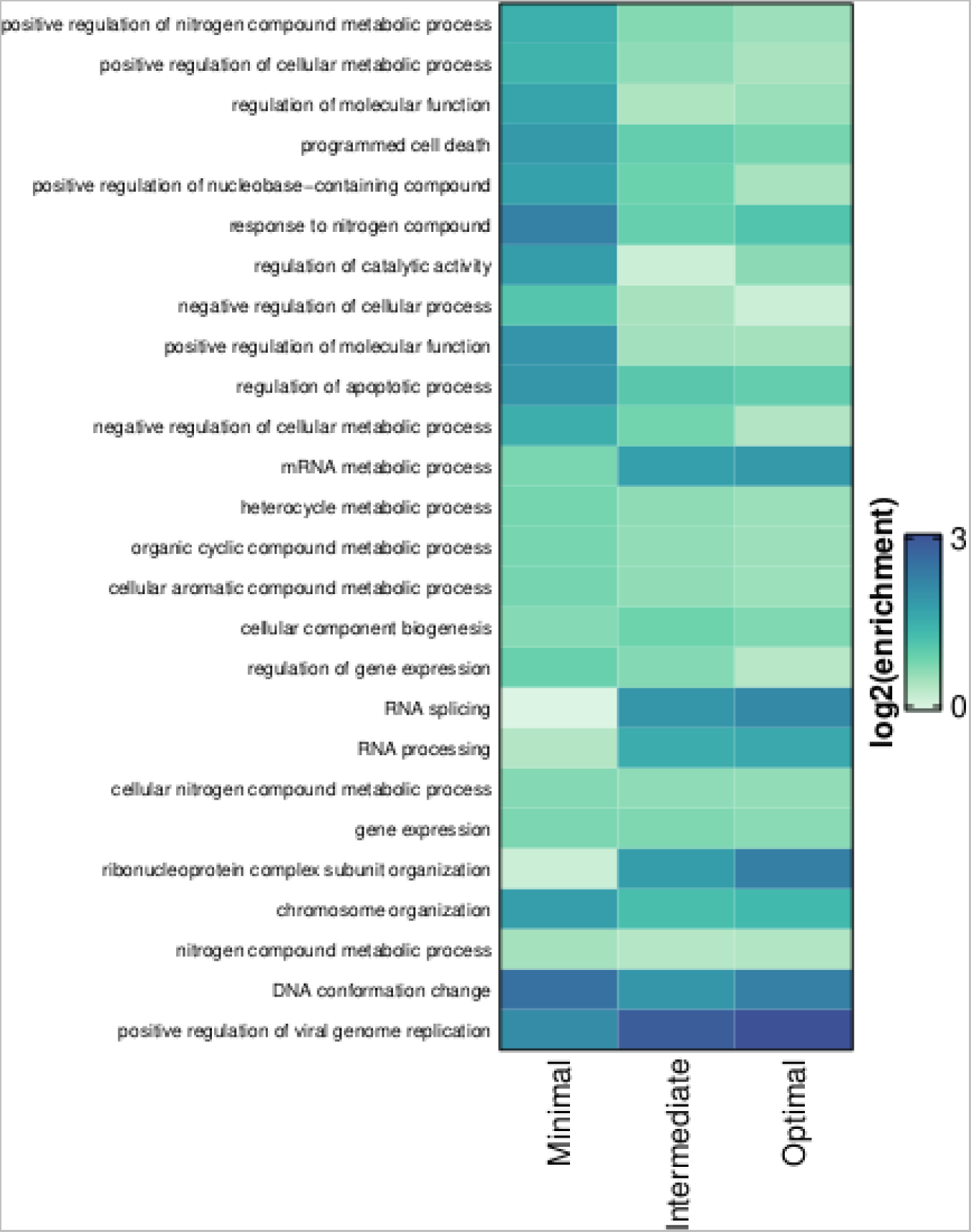
GO term enrichment analysis of human CK2 substrates. Gene ontology (GO) term enrichment analysis of minimal (SQ < 0.3), intermediate (0.3 < SQ < 0.6), and optimal (SQ > 0.6) CK2 sites for biological process (BP) terms. For each category (minimal, intermediate, optimal), the top 20 GO terms were selected and then conceptually redundant GO terms (e.g. RNA processing and mRNA processing) were filtered. Terms across each category were combined and then the enrichment ratios (foreground / background) calculated. SQ: substrate quality.

**Supplementary figure 11 -.**
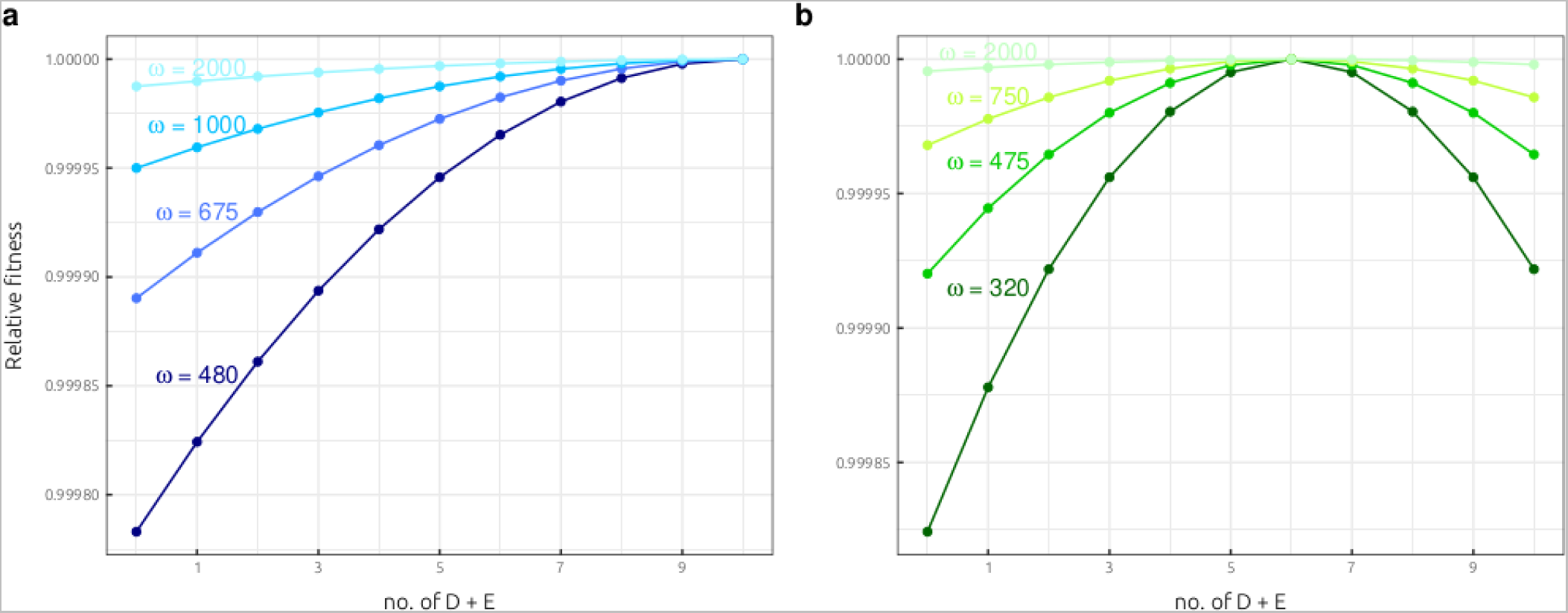
Examining different fitness functions for the CK2 sequential-fixation model. Gaussian fitness functions for the sequential-fixation model of CK2 substrate evolution presented in Figure 3. Panel **a)** corresponds to directional selection (θ=10) whereas panel **b)** corresponds to stabilising selection (θ=6). Changing the scale parameter (ω) modulates the relative strength of selection between genotypic states.

**Supplementary figure 12 -.**
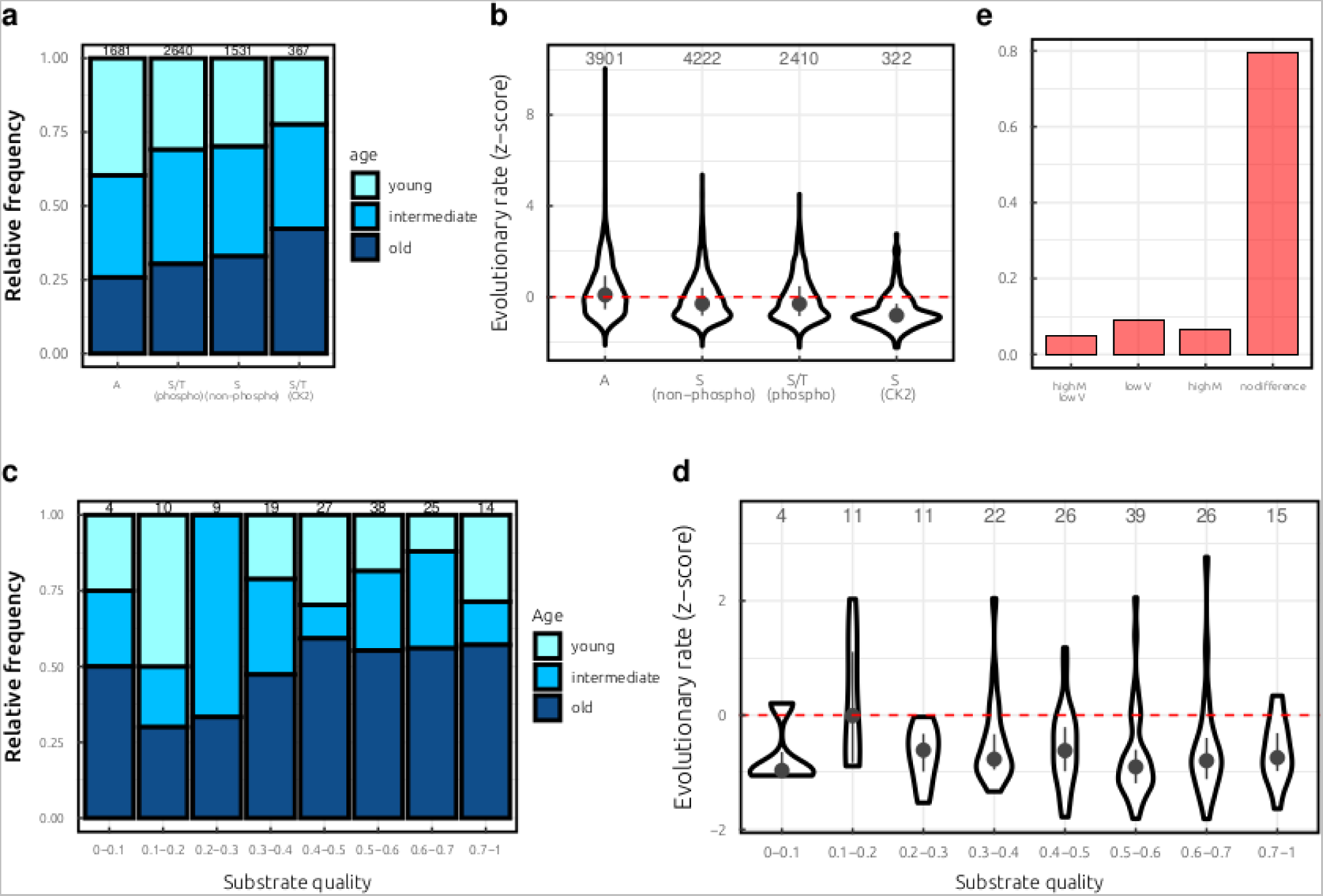
The evolution of CK2 target sites across vertebrate species. **a)** Estimated age of random A residues, phosphorylated S/T, non-phosphorylated S, and CK2 target phosphosites, all within known CK2 substrates. Age was estimated conservatively from the ancestral sequence reconstructions by identifying the earliest (closest to the root) phosphorylatable residue in the phospho-acceptor position. For random A residues, the earliest (closest to the root) A residue in the ancestral sequence root was considered. young: <200 mya, intermediate: 200<x<600 mya, old : >600 mya. Mya: million years ago **b)** Evolutionary rate of random A residues, non-phosphorylated S, phosphorylated S/T, and CK2 target phosphosites, all within known CK2 substrates. Evolutionary rate z-score normalised per-protein within disordered regions. **c)** Estimated age of CK2 substrates with respect to substrate quality for high-confidence multiple sequence alignments (MSAs). Age was estimated conservatively from the ancestral sequence reconstructions by identifying the earliest (closest to the root) phosphorylatable residue in the phospho-acceptor position. young: <200 mya, intermediate: 200<x<600 mya, old : >600 mya. Mya: million years ago **d)** Relationship between substrate quality and evolutionary rate for high-confidence MSAs. Evolutionary rate z-score normalised per protein within disordered regions. **e)** Summary across 141 CK2 substrates of phosphosites with a significantly higher median (M) and lower variance (V) than the simulated sequences **(n=7)**, a significantly lower variance **(n=13)**, a significantly higher median **(n=9)**, or no significant difference detected **(n=112).** For this analysis, only CK2 substrate sites with D/E+1 and D/E+3 were considered, and substrate positions +1 and +3 were not considered for the substrate quality calculations. The y-axis corresponds to the relative frequency of CK2 substrates (between 0 and 1).

**Supplementary figure 13 -.**
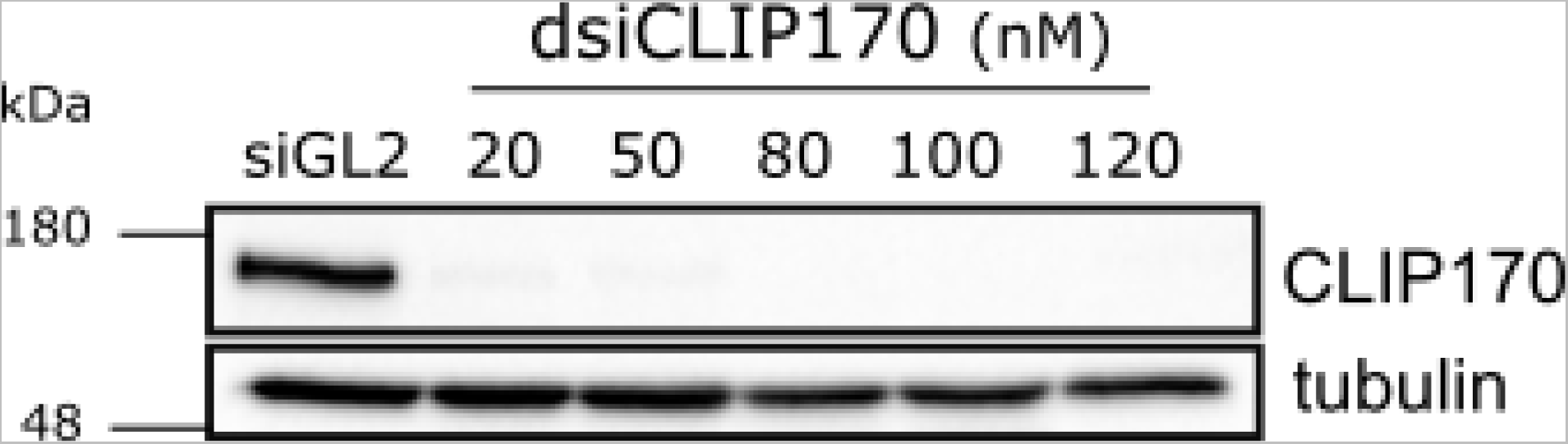
Depletion of endogenous CLIP-170 using dsiRNA. HeLa T-REx cells were treated with control (siGL2) or CLIP170 (dsiCLIP170) dsiRNAs at the indicated concentrations. CLIP170 protein is in the upper panel. Tubulin (bottom panel) illustrates equal loading.

## Methods

### Specificity modelling and stoichiometry analysis

Kinase-substrate relationship (KSR) data was retrieved from the ProtMapper resource (Bachman, Gyori, and Sorger 2022), which compiles kinase substrates from databases such as PhosphoSitePlus and Signor (Hornbeck et al. 2019; Licata et al. 2020). We exclude kinase-substrate annotations supported only by text mining tools, which may reflect indirect interactions (Bachman, Gyori, and Sorger 2022). For CK2 substrates, we exclude as potential confounders CK2 targets with an S at positions +1 and/or +3, which could indicate CK2 hierarchical phosphorylation sites (N. St-Denis et al. 2015). The preference of human CK2 for D/E at each substrate position **(Supplementary figure 1a)** was assessed with the online pLogo tool using default parameters and the human proteome as a background (O’Shea et al. 2013).

Calculation of substrate quality for CK2 target sites is derived from the CK2 position weight matrix (PWM) constructed simply on the relative amino acid frequencies for substrate positions −6 to +6. Relative amino acid frequencies for D and E (combined) are summed across each D/E-containing position in the peptide, and then divided by the maximum possible score i.e. the summed relative D/E frequencies across all positions (−6 to +6). For the analysis in **Supplementary figure 1b**, the same score is calculated but instead using PWM scores from a previous peptide array study performed on CK2 (Hutti et al. 2004). In **Supplementary figure 3a and 3b**, full 20 × 13 PWMs are derived from literature data (Bachman, Gyori, and Sorger 2022) and MS/MS data (Sugiyama, Imamura, and Ishihama 2019) using a method described previously (Invergo et al. 2020) while also controlling for the disorder-biased amino acid composition in the background proteome using data from (Dubreuil, Matalon, and Levy 2019). In **Supplementary figure 3c**, the peptide array PWM was sourced directly from (Johnson et al. 2023).

CK2-specific stoichiometry data (PC9 cells) was retrieved from a previous study (Tsai et al. 2015). Stoichiometry values were taken only for singly-phosphorylated peptides. We also excluded potential hierarchical phosphorylations sites (i.e with S at +1 and/or +3) and for the **Figure 1** and **Supplementary figure 1** analysis only serine-phosphorylated peptides were retained as the phosphoacceptor residue also has a strong impact on stoichiometry **(Supplementary figure 5e)**. EGFR- and MAPK-specific stoichiometry data were taken from the same study and filtered to include singly phosphorylated peptides only. For the EGFR analysis, any site with D/E at −4, or D/E at −3, or D/E/F at +2, or I/P/L/M at +3 was considered ‘not minimal’, on the basis of a previous deep sequencing-based assay (Cantor, Shah, and Kuriyan 2018).

Kinase-agnostic stoichiometry data was taken from three sources (Olsen et al. 2010; Wu et al. 2011; Sharma et al. 2014). As with the CK2-specific data, we retain only singly phosphorylated peptides with a serine phosphoacceptor. Where this information is available, we also retained Class I phosphorylation sites (localisation probability > 0.75) only (Olsen et al. 2006; Sharma et al. 2014). CK2 substrates were predicted using the **S-D/E-x-D/E** motif. For the Olsen *et al* data the maximum stoichiometry across the six cell cycle stages was taken. For the Sharma *et al* data, stoichiometry values were taken for the control condition (asynchronous cells).

### Generalised linear models of stoichiometry

For the GLM (generalised linear model) analysis of phosphorylation stoichiometry, the CK2 stoichiometry data was retrieved and processed as described directly above. The response variable (stoichiometry) is in the range of 0 to 1 and was modelled using a quasi-binomial family GLM with a *cloglog* link function. All GLMs were generated using the *glm()* function in R, and comparisons between different models were made with an F-test in R using the *anova(m1, m2, test=’F’)* function.

The significance of any given explanatory variable (substrate quality, abundance, etc.) was assessed by comparing a minimal model (with one explanatory variable) against a null model without any explanatory variables, using the *anova()* function in R e.g. *anova(m_null_, m_minimal_, test=’F’).* The same GLM methodology was used for all CK2, MAPK, and EGFR GLMs.

Substrate disorder was predicted using an AlphaFold2 (AF2)-based method in which the smoothened relative surfaces accessibility (RSA) in a 15 amino acid window was used as a proxy for protein disorder (Akdel et al. 2022). Phosphosites with a smoothened RSA greater than or equal to a cut-off of 0.55 were considered to be disordered. Proteins without an AF2 model were predicted using the *SPOT-Disorder-Single* tool (Hanson, Paliwal, and Zhou 2018). Protein abundances for substrates were retrieved from the *PaxDB* database (M. Wang et al. 2012). Subcellular location data was retrieved from UniProtKB by extracting GO terms in the ‘cellular compartment’ category (UniProt Consortium 2021). On this basis, substrates were assigned either to nuclear (N), cytoplasmic/cytosolic (C), combined nuclear:cytoplasmic (NC), or ‘Other’ categories if the protein was found in a different organelle (e.g. mitochondria). Protein-protein interaction (PPI) data was retrieved from *iRefIndex*, which collates PPIs from several databases (Razick, Magklaras, and Donaldson 2008).

### Phosphosite functional scores

Phosphosite functional scores **(Figure 2a** and **Supplementary figure 6a)** were derived from the *funscoR* package introduced in (Ochoa et al. 2019). Training was performed on the ‘reference phosphoproteome’ of 116,258 non-redundant phosphosites in human (Ochoa et al. 2019). We tested the relationship between substrate quality and the functional scores in **Figure 2a** and **Supplementary figure 6a**; therefore, features related to kinase specificity (’isELMLinearMotif’, ‘isMotif’, ‘PWM_max_mss’, ‘netpho_max_KIN’, and ‘netpho_max_all’) were excluded from the dataset before model training to prevent circularity between model training and model analysis. In **Supplementary figure 6a**, phosphosites scores are assigned to predicted CK2 targets. These correspond to *S-D/E-x-D/E* phosphosites found on proteins that are in close physical proximity to human CK2 (on the basis of BioID data) (Niinae et al. 2021)

### Phosphoproteome analysis

For the **Figure 2b** and **Supplementary figure 7** analysis, phosphoproteome data across 435 conditions were obtained from the http://phosfate.com/ resource corresponding to the (Ochoa et al. 2016) publication. Quantifications for individual phosphopeptides were extracted from the MySQL database of http://phosfate.com/ using only sites with a localisation probability > 0.75. As in **Supplementary figure 5a**, the abundance data for human proteins (whole organism - integrated) used for the analysis in **Supplementary figure 7a** and **Supplementary figure 7b** were obtained from *PaxDB* (M. Wang et al. 2012). ‘High’ abundance proteins shown in **Figure 2a** correspond to a ppm value greater than 100 parts per million (ppm), representing proteins in the top 6.6% of abundance values for human. For the analysis of phosphosite abundance **(Supplementary figure 7a-b)**, phosphosites of known stoichiometry were extracted from (Sharma et al. 2014). Only singly-phosphorylated sites with a localisation probability > 0.75 were retained. The phosphosite abundance was calculated by multiplying the average stoichiometry (under the control condition) by the protein abundance reported in PaxDB. The ion mobility of phosphopeptides (**Supplementary figure 7c** and **7d**) was predicted with AlphaPeptDeep using default parameters (Zeng et al. 2022).

For the **Figure 2c** analysis, yeast phosphoproteome data across *~*100 conditions were retrieved from (Leutert et al. 2022), using all conditions where phosphoproteome changes were measured in 5 minutes or less. Known CK2 substrates in *S. cerevisiae* were compiled from BioGRID and a manual curation of the literature (Oughtred et al. 2021). Predicted substrates were made using interactors from BioGRID and YeastKID (Sharifpoor et al. 2011), and matching phosphosites to the S-D/E-x-D/E motif. We consider a phosphosite from the core yeast phosphoproteome to be regulated if it has a log2-fold increase (relative to the control) of at least 1 (−1 for downregulated) and an adjusted p-value < 0.05.

The cell cycle phosphoproteome data in **Figures 2d, 2e**, and **Supplementary figure 9** derive from a previous study where the phosphoproteome was sampled in human cells at six cell cycle stages (Olsen et al. 2010). We filtered these data to retain only singly phosphorylated peptides corresponding to Class I phosphosites. In addition, phosphosites with missing values in any one of the six stages were removed from the analysis. Only known CK2 substrates were retained for the analysis in **Figures 2d and 2e** and predicted CK2 targets (on the basis of p*S-D/E-x-D/E* phosphosites and BioID data) were selected for the analysis in **Supplementary figures 9a** (Olsen et al. 2010; Niinae et al. 2021). For a given phosphosite, the number of additional upstream kinases in **Supplementary figures 6b and Supplementary figures 9b** were calculated from the *ProtMapper* dataset of kinase-substrate relationships described above (Bachman, Gyori, and Sorger 2022).

### GO term enrichment

The gene ontology (GO) term enrichment analysis presented in **Figure 2f** and **Supplementary figure 10** was performed in R using the t*opGO* library (Alexa and Rahnenfuhrer 2016). We take as our background for the GO term enrichment all proteins with at least one high-confidence phosphosite recorded in (Ochoa et al. 2019). As the foreground we take all known and predicted CK2 substrates, again making predictions on the basis of CK2 *S-D/E-x-D/E* motif matches to CK2 transient interactors (Olsen et al. 2010; Niinae et al. 2021). GO enrichment was performed for ‘biological process’ terms using the classic algorithm (‘classCount’) and Fisher’s exact test (‘GOFisherTest’) test statistic (Alexa and Rahnenfuhrer 2016).

### Sequential-fixation model for CK2 substrate evolution

The sequential-fixation model presented in **Figure 3** was first presented in general form by Lynch and colleagues (Lynch and Hagner 2015; Lynch and Trickovic 2020) and in more detail by Lynch in (Lynch 2020). We describe here how we apply this model to the evolution of CK2 target sites. As described in the *Results* section, we subset the kinase-substrate relationships (KSRs) for all known human CK2 substrates with aspartate (D) or glutamate (E) at positions +1 and +3 exclusively, and then calculate the D/E distribution for *all other* positions from −6 to +6.

We first present the results of the model under selective neutrality. Mutations to D and E are considered as positive (+) alleles because they make a positive contribution to the trait (i.e. they increase substrate quality). Conversely, mutations from D or E to any other amino acid are considered to be a negative (−) allele because they make a negative contribution to the trait (i.e. they decrease substrate quality) **(Figure 3a)**. The probability of genotypic states (D/E = 0, D/E = 1, D/E = 2) is then given by the binomial distribution:

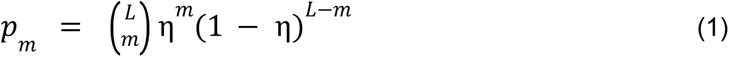

Where *L* is the number of mutable sites contributing towards a trait (here *L* = 10), m is the number of sites with a + allelic state (here a D or E), *p_m_* is the equilibrium probability of each genotypic state (here the relative frequency of sites with 0 D/E, 1 D/E, 2 D/E, etc). η corresponds to the expected background frequency of + alleles (D or E), and is given by:

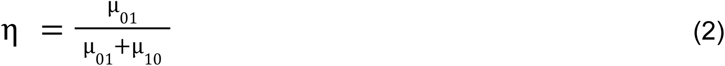

Where *μ*_01_ is the ‘forward’ mutation rate to D/E residues and *μ*_10_ is the ‘reverse’ mutation rate away from D/E. Using per-nucleotide mutation rates estimated for humans and knowledge of the genetic code gives an estimated η of **6.74%** (Carlson et al. 2018). Simply multiplying human nucleotide frequencies for each D/E-encoding codon and then summing the result across codons gives a similar estimated η of **6.86%** (Gardini et al. 2016). However, the observed frequency of D/E in the proteome is known to deviate far from the neutral expectation (Gardini et al. 2016), possibly because of the role of D and E as disorder-promoting amino acids. Given that most phosphosites are found in disordered regions (Iakoucheva et al. 2004), we used disordered content predictions across the human proteome (Dubreuil, Matalon, and Levy 2019) to predict the expected frequency of D/E in random phosphosites (kinase-agnostic) while controlling for the bias of S/T phosphosites towards disordered regions. This gives a new predicted η of **13.75%** that we use to produce the results in **Figure 3b**.

In **Figure 3c** and **3d** the model is updated to account for the role of selection. Under selection, the rate of transition to a ‘higher’ genotypic state (i.e. with more D/Es) is proportional to the product of the forward rate of mutation (*μ*_10_) and the forward probability of fixation (φ_10_), and *vice versa (Lynch and Trickovic 2020)*. The equilibrium frequency of each state is equal to the product of the forward and backward rates, which simplifies to the following equation under selection (Lynch and Trickovic 2020):

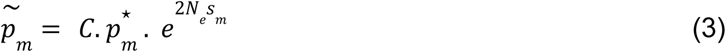

Where 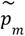 corresponds to the equilibrium genotype frequencies under selection and 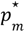 to the frequencies under neutrality, as set out in equation **(1)**. *C* is a normalisation constant that ensures all of the probabilities sum to 1. *N_e_* is the human effective population size **(2.1×10^4^)** and is taken from (Lynch and Trickovic 2020), as was the *N_e_* for *S. cerevisiae*. s*_m_*is a fitness parameter that represent the fitness loss relative to the fitness optimum (set at 1):

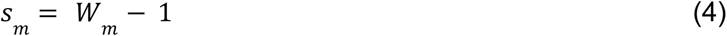

The fitness (*W_m_*) for any genotypic value of *m* is defined by a fitness function. Here we specify a Gaussian fitness function **(Supplementary figure 11),** as in (Lynch and Trickovic 2020) and (Lynch 2020):

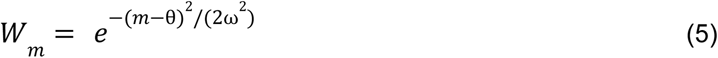

As described in the *Results, θ* is a location parameter that specifies the fitness optimum whereas ω is a ‘scale’ parameter that specifies how narrow or broad the fitness function is **(Supplementary figure 11)**. *θ a*nd ω are unknown parameters for real CK2 sites, but in **Figure 3d** the parameter space for *θ a*nd ω is sampled systematically and the model outputs are compared to the empirical distribution using the Kulback-Leibler divergence (Kullback and Leibler 1951). *θ* was sampled from 0 to 10 and ω sampled from 20 to 1200 in steps of 20.

For the analysis of yeast CK2 substrates in **Figure 3e**, known substrates **(Supplementary Table 5)** were compiled from BioGRID and a manual curation of the literature (Oughtred et al. 2021).

### Conservation analysis

One-to-one vertebrate orthologues of CK2 substrates were retrieved from Ensembl Compara using the Rest API (November 2021) (Herrero et al. 2016). Predicted hierarchical phosphorylation sites were excluded from the analysis (N. St-Denis et al. 2015), as described above. We restricted the analysis to CK2 target sites mapping to disordered regions given that the analysis of ordered regions may be confounded by structural constraints of the protein domain, and that ordered and disordered regions tend to evolve at different rates on average (Brown et al. 2002; Brown, Johnson, and Daughdrill 2010; Langstein-Skora et al. 2022). Disordered regions were predicted using an AlphaFold2-based method, as described above (Akdel et al. 2022). Orthologous sequences were aligned using the L-INS-i method of MAFFT (Katoh et al. 2005). Predicted spurious orthologous sequences were removed using trimAl (*-resoverlap 0.75 -seqoverlap 80*) (Capella-Gutiérrez, Silla-Martínez, and Gabaldón 2009). Phylogenies were generated using FastTree with default parameters (Price, Dehal, and Arkin 2010). Taxonomic annotations for each species given in Ensembl were used to automatically root the tree by selecting the oldest clade (species in taxa most distantly related to the *Homininae*) as an outgroup.

The estimates of phosphosite ages were based upon using ancestral sequence reconstruction to identify the earliest (i.e. closest to the root) ancestral node that is phosphorylatable (contains an S or T). This is a conservative and approximate method since phosphorylation may have evolved some time after the appearance of a suitable phosphoacceptor (Landry, Levy, and Michnick 2009). Ancestral sequence reconstruction was performed using FastML, employing a maximum likelihood-based approach for the marginal reconstruction of substitutions and indels (Ashkenazy et al. 2012). FastML was executed using a disorder-based substitution matrix given that we restrict our evolutionary analysis to disordered phosphosites (Szalkowski and Anisimova 2011). Approximate ages for ancestral nodes were predicted using the Ensembl taxonomy annotations combined with taxon divergence estimates given in TimeTree (Kumar et al. 2017).

Evolutionary rate estimates were generated by running the rate4site software with a Gamma model-based optimisation of branch lengths (Mayrose et al. 2004). As an input we use the MSA and subtree corresponding to sequences with an ancestral residue S or T at the phospho-acceptor position, based upon ancestral sequence reconstructions **(Figure 4a).** Z-score normalisation was then performed for each phosphosite by considering only predicted disordered regions in the alignment. CK2 substrates with fewer than 20 predicted disordered residues were removed. Phosphorylated residues labelled in **Supplementary figure 12a** and **Supplementary figure 12b** correspond to high-confidence phosphosites from (Ochoa et al. 2019) mapping to CK2 substrates.

Controlling for alignment quality **(Supplementary figure 12c** and **Supplementary figure 12d)** was performed using a similar approach as in (Studer et al. 2016). Specifically, phosphosite positions with more than 20% gaps were excluded, as were phosphosites where more than 20% of orthologues contain at least one S residue in a window from positions −3 to +3.

### Simulated evolution analysis

Simulated evolution was performed 200 times for each set of orthologous sequences using Pyvolve (Spielman and Wilke 2015). As illustrated in **Figure 4a**, the simulations take as an input the earliest (closest to the root) phosphorylatable sequences predicted by ancestral sequence reconstruction, and then simulations are performed along all descendant branches in the phylogeny. The simulations were executed using a disorder-based substitution matrix (Szalkowski and Anisimova 2011), and disorder-controlled equilibrium amino acid frequencies calculated using the approach described in the above section (*Sequential-fixation model for CK2 substrate evolution) (Dubreuil, Matalon, and Levy 2019)*. The simulations were run separately for each substrate position (−6 to +6), each time scaling the branch lengths by the position-specific evolutionary rates calculated by rate4site (Mayrose et al. 2004). Outputs for each substrate position (−6 to +6) were then concatenated (per trial) to generate the simulated substrate 13-mer.

### Peptide array analysis

The phosphorylation intensity of CK2 substrates represented in **Figure 1d**, **Figure 5** and **Supplementary figure 2-3** were assayed using peptide arrays.

The CK2 peptide array synthesis was performed by the Francis Crick Institute Peptide Chemistry Science Technology Platform. The peptide array was synthesised on an Intavis ResResSL automated peptide synthesiser (Intavis Bioanalytical Instrument, Germany) by cycles of N(a)-Fmoc amino acids coupling via activation of the carboxylic acid groups with diisopropylcarbodiimide in the presence of ethylciano-(hydroxyamino)-acetate (Oxyma Pure) followed by removal of the temporary-amino protecting group by piperidine treatment. Subsequent to chain assembly, side chain protection groups are removed by treatment of the membrane with a deprotection cocktail (20 ml 95% trifluoroacetic acid, 3% triisopropylsilane and 2% H_2_O) for 4 hours at room temperature, then washing (4x dichloromethane, 4x ethanol, 2x H_2_O and 1x ethanol) prior to being air dried. The final product was a cellulose membrane containing 748 13-mers peptides including one positive control (RRRAEESEDEEEA). The full list of peptides can be found in **Supplementary table 7**.

The cellulose membrane was placed in an incubation trough and moistened with 10 ml ethanol. It was subsequently washed twice with 50 ml kinase buffer (10 mM Tris-HCl, pH 7.4, 50 mM KCl, and 10 mM MgCl_2_ (Hutti et al. 2004) and incubated overnight in 100 ml reaction buffer (kinase buffer + 0.2 mg/ml BSA (BSA Fraction V, Sigma) + 50 µg/ml kanamycin). The next day, the kinase buffer was removed and the membrane was incubated at 30°C for 1 hour in 30 ml blocking buffer (kinase buffer + 1 mg/ml BSA + 50 µg/ml kanamycin). After incubation, the blocking buffer was replaced with 30 ml reaction buffer supplemented with 300 µl 10 mM ATP and 125 µCi [^32^P]-ATP (Hartmann Analytics, Germany). The reaction was started by adding 5000 units of recombinant Casein Kinase II (NEB, P6010S) and left to incubate for 20 min at 30°C with gentle agitation. After incubation, the reaction buffer was removed and the membrane was washed 10 × 15 min with 100 ml 1M NaCl, 3 × 5 min with 100 ml H_2_O, 3 × 10 min with 5% H_3_PO_4_, 3 × 5 min with 100 ml H_2_O and 2 × 2 min with 100 ml ethanol. The membrane was left to air dry before being wrapped up in plastic film and exposed overnight to a PhosphorScreen. The radioactivity incorporated into each peptide was then determined using a Typhoon FLA 9500 phosphorimager (GE Healthcare).

For the CLIP-170 S1364 phosphosite, vertebrate one-to-one and one-to-many orthologues were retrieved using Ensembl Compara (June 2021) (Herrero et al. 2016). Unique 13-mer (−6 to +6) peptides were taken forward for the peptide array analysis. Mutants to systematically decrease and increase CLIP-170 S1364 substrate quality **(Figure 5e-f)** were designed manually. The CLIP-170 full-length sequences were aligned using the L-INS-i method of MAFFT and then manually trimmed (Katoh et al. 2005). The CLIP-170 phylogeny was generated with IQ-TREE2, using an automated approach to identify the best substitution model (L.-T. Nguyen et al. 2015; Kalyaanamoorthy et al. 2017). All CLIP-170 S1364 mutant and orthologue peptide sequences are given in **Supplementary table 7** alongside their measured phosphorylation values.

### CLIP-170 vector construction

To assemble the CLIP-170 sequence into the pcDNA5/FRT/TO-3xMyc-eGFP vector, the pcDNA5/FRT/TO-3xMyc-eGFP plasmid was first doubly digested (2 µg of DNA digested sequentially with 80 units of BamHI for 1h30 and with 20 units of EcoRV for 1h30). Then, four fragments covering the entire sequence of CLIP-170 (splice isoform NM_198240.3; Fragment 1 (4-1295; without the ATG), Fragment 2 (1296-2587), Fragment 3 (2588-3879) and Fragment 4 (3880-4179)) were synthesised by Twist Bioscience (USA). Silent mutations were introduced in Fragment 1 for amino acids at positions between 251 and 257 (GGAAGAATGATGGCGCTGTT changed to GcAAGAAcGAcGGgGCcGTg) so that the construct would not be targeted by the siRNA used to repress the endogenous copy. In total, five different versions of Fragment 4 (WT and four mutants) were designed, one for each mutant needed. Synthesised Fragments 1 to 3 were amplified with primers CLOP267-D5/E5, CLOP267-F5/G5 and CLOP267-H5/A6 respectively (PCR program: 5 min at 95 °C, 5 cycles: 20 s at 98 °C, 15 s at 60 °C, 1 min at 72 °C, 30 cycles: 20 s at 98 °C, 1 min 15 s at at 72 °C, and a final extension of 3 min at 72 °C) while the different versions of Fragments 4 were amplified using CLOP267-B6/CLOP273-A8 (PCR program: 5 min at 95 °C, 5 cycles: 20 s at 98 °C, 15 s at 64 °C, 20 s at 72 °C, 30 cycles: 20 s at 98 °C, 35 s at at 72 °C, and a final extension of 3 min at 72 °C). In these PCR amplifications, overlapping sequences of 20 bp were added at the extremities to allow Gibson assembly in the vector. The digested vector along with PCR-amplified fragments were all purified on magnetic beads (Axygen AxyPrep Mag PCR Clean-up Kit) before being assembled by Gibson DNA assembly (Gibson et al. 2009), producing plasmids pcDNA5/FRT/TO-3xMyc-eGFP-CLIP-170WT and its different mutants. These plasmids were retrieved by transformation in MC1061 homemade chemically competent bacteria (Green and Rogers 2013)

Even if all assemblies were successful, unfortunately, only pcDNA5/FRT/TO-3xMyc-eGFP-CLIP170mut2 was found to be without any point mutation (full-length plasmid sequencing, Plasmidsaurus, SNPsaurus, USA). Starting from pcDNA5/FRT/TO-3xMyc-eGFP-CLIP170mut2, we thus reintroduced the correct mutations by site-directed mutagenesis based on the QuickChange™ Site-Directed Mutagenesis System (Stratagene, La Jolla, CA) to recreate the WT and the three other mutants. Briefly, we amplified the pcDNA5/FRT/TO-3xMyc-eGFP-CLIP170mut2 plasmid using pairs of primers containing the desired mutation at the centre (see **Supplementary table 9** for the primers specific to each mutation) (PCR program: 5 min at 95 °C, 22 cycles: 20 s at 98 °C, 15 s at 68 °C, 10 min at 72 °C, and a final extension of 10 min at 72 °C). The PCR products were then incubated for 1 h 30 at 37 °C with 6 U of DpnI enzyme to remove parental DNA and mutated plasmids were retrieved directly by transformation in MC1061 homemade chemical competent bacteria. The mutation insertions were confirmed by Sanger sequencing (Plateforme de séquençage et de génotypage des génomes, CRCHUL, Canada) using CLOP267-H6 as a sequencing primer.

All PCR reactions mentioned above were performed with oligonucleotides defined in **(Supplementary table 9)**, using KAPA HiFi HotStart polymerase.

### Generation of stable cell lines

Stable cell lines were generated using Flp-In T-REx system (Life Technologies). Transfection was done with FuGENE® HD (Promega) reagent according to the manufacturer’s instructions. The transfection reagent was mixed with the pOG44 Flp-Recombinase expression vector and the pCDNA5/TO/FRT-3xMyc-eGFP-CLIP-170 WT or one of the mutant vectors with a 3:1 ratio in Opti-MEM^TM^ I medium (Gibco, ThermoFisher Scientific). Selection started 48h after the transfection in a Dulbecco’s modified Eagle medium (DMEM, Sigma) supplemented with 10% of bovine growth serum (Cytiva), 100 U/ml penicillin and 100 µg/ml streptomycin (MultiCell, Wisent) and in the presence of hygromycin B (200 µg/ml, MultiCell, Wisent) and blasticidin (10 µg/ml, InVivoGen) at 37°C and 5% CO_2_. Media was replaced three times a week for three weeks and then four pools were generated by vectors and grown separately. Expression of the CLIP-170 constructs was then tested by IF after 24h of doxycycline treatment (1000 ng/ml, TCI America) and the pool with the best 3xMyc-eGFP-CLIP-170 expression was kept for the further experiment.

### Immunofluorescence

1,5×10^5^ Cells were seeded on 15 mm round glass coverslips coated with polyethylenimine (PEI) solution (25 g/ml PEI, Polysciences Inc. and 150 nM NaCl) for 48 hours. Subsequently, 80 nM CLIP1 (CLIP-170) dsiRNA (target region 752 to 771 aa : 5’-GGAAGAATGATGGCGCTGTT-3’) were transfected using jetPrime® (Polyplus) for 48 hours for endogenous CLIP-170 knock-down. Twenty-four hours before fixation, the cells were induced with doxycycline (1000 ng/ml) followed by a synchronisation in mitosis with 330 nM of nocodazole (Sigma-Aldrich) for the last 16 hours. Cells were pre-permeabilized for 60 seconds with 0.1% Triton X-100 in PEM (20 mM PIPES pH 6.9, 0.2% Triton X-100, 10 mM EGTA pH8.0, 1 mM MgCl_2_) followed by fixation (3,7% formaldehyde in PEM) for 5 minutes (Tanenbaum et al. 2006). After fixation, cells were blocked in 3% BSA in PBS with 0,2% Tween 20 and incubated overnight with anti-MYC (A-14, Santa Cruz Biotechnology, 2 ug/ml), anti-CENP-C (PD030, MBL International, 1 ug/ml) and anti-CENP-A (3-19, Enzo Life Sciences, 1 ug/ml) at 4°C in a humid chamber. AlexaFluor-AffiniPure series secondary antibodies (Jackson ImmunoResearch) and Hoechst 33342 (ThermoFisher, 1 mg/ml stock) were used at 1:1000 and incubated for 2h at room temperature.

Immunofluorescence images were acquired with an Olympus IX80 inverted confocal microscope equipped with a WaveFX-Borealin-SC Yokagawa spinning disc (Quorum Technologies) and an Orca Flash4.0 camera (Hamamatsu). Image acquisition was performed using Metamorph software (Molecular Devices). Optical sections were acquired with identical exposure times for each channel within an experiment and then projected with maximum intensity into a single picture using ImageJ2 (Fiji version 1.53t) (Rueden et al. 2017; Schindelin et al. 2012). The kinetochore signal intensity was measured according to an established protocol (Saurin and Kops 2016) using ImageJ2. Briefly, cells were measured individually by cropping the desired cell, creating a selection of the kinetochores area by thresholding (CENP-C signal), and transferring to and measuring the selection in the desired channels. CLIP-170 intensity was normalised relative to the CENP-C signal for each cell. For image processing, Quickfigure (ImageJ PlugIn, (Mazo 2021) was used and all images shown in the figure have been identically scaled, brightness and contrast adjusted at the same value for the same channel and are representative of the results obtained.

## Data availability

The code generated for this analysis is available at: https://github.com/Landrylab/Bradley_2023_CK2

## Acknowledgements

We would like to thank Stephanie J Spielman for assistance with the evolutionary simulations. We would also like to thank the Landry lab for their helpful comments throughout the project. DB was funded by the European Molecular Biology Organization (EMBO) Long-Term Fellowship (ALTF 1069-2019). HB and MT are supported from the Francis Crick Institute which receives its core funding from Cancer Research UK (CC2132), the UK Medical Research Council (CC2132), and the Wellcome Trust (CC2132). We thank Nicola O’Reilly for generating the peptide arrays. The peptide chemistry platform at the Francis Crick Institute receives funding from Cancer Research UK (CC0199), The UK Medical Research Council (CC0199) and the Wellcome Trust (CC0199). For the purpose of Open Access, the authors have applied a CC BY public copyright licence to any Author Accepted Manuscript version arising from this submission. SE is funded by grants from the Canadian Institutes of Health Research (RN376557 RN486735) and the National Science and engineering council of Canada (RGPIN-2016-05841). The project was funded by a HFSP grant RGP0034/2018 and by a Canadian Institutes of Health Research Foundation grant number 387697 to CRL.

## Author contributions

CRL and DB conceived the project. DB performed all computational analysis. CRL supervised the project. DB wrote the manuscript. CRL, DB, IGA, MT and SE reviewed and edited the draft manuscript. CRL, DB, MT, and SE designed the experiments. HB performed the peptide array analysis. IGA generated the mutants for CLIP-170. CG performed the *in vivo* analysis.

